# The protective function of an immunity protein against the *cis*-toxic effects of a *Xanthomonas* Type IV Secretion System Effector

**DOI:** 10.1101/2023.08.28.555164

**Authors:** Gabriel U. Oka, Diorge P. Souza, Germán G. Sgro, Cristiane R. Guzzo, German Dunger, Chuck S. Farah

**Affiliations:** Departamento de Bioquímica, Instituto de Química, Universidade de São Paulo, São Paulo, SP, Brazil; Structure and Function of Bacterial Nanomachines, Institut Européen de Chimie et Biologie – CNRS, UMR 5234 Microbiologie Fondamentale et Pathogénicité University of Bordeaux, Pessac, France; Division of Cell Biology, MRC Laboratory of Molecular Biology, Cambridge, United Kingdom; Departamento de Ciências BioMoleculares, Faculdade de Ciências Farmacêuticas de Ribeirão Preto, Universidade de São Paulo, Ribeirão Preto, SP, Brazil; Departamento de Microbiologia, Instituto de Ciências Biomédicas, Universidade de São Paulo, São Paulo, SP, Brazil; Instituto de Ciencias Agropecuarias del Litoral (ICiAgro Litoral), Universidad Nacional del Litoral, CONICET, Facultad de Ciencias Agrarias, Esperanza, Argentina

## Abstract

Many bacterial species use specialized secretion systems to translocate proteinaceous toxic effectors into target bacterial cells. In most cases, effectors are encoded in bicistronic operons with their cognate immunity proteins. The current model is that immunity proteins could, in principle, provide protection in two different ways: i) by avoiding self-intoxication (suicide or *cis*-intoxication) or ii) by inhibiting intoxication due to “friendly-fire” translocation from neighboring sister cells (fratricide or *trans*-intoxication). Here, we set out to distinguish between these two protection mechanisms in the case of the bactericidal *Xanthomonas citri* Type IV Secretion System (X-T4SS), where killing is due to the action of a cocktail of secreted effectors (X-Tfes) that are inhibited by their cognate immunity proteins (X-Tfis). We use a set of *X. citri* mutants lacking multiple X-Tfe/X-Tfi pairs to show that X-Tfis are not absolutely required to protect against *trans*-intoxication. Our investigation then focused on the *in vivo* function of the lysozyme-like effector X-Tfe^XAC2609^ and its cognate immunity protein X-Tfi^XAC2610^. We observe the accumulation of damage in the *X. citri* cell envelope and inhibition of biofilm formation due to the action of X-Tfe^XAC2609^ in the absence of X-Tfi^XAC2610^. We show that X-Tfe^XAC2609^ toxicity is independent of an active X-T4SS and that X-Tfi^XAC2610^ protects the cell colony against X-Tfe^XAC2609^-induced *cis*-intoxication via autolysis. *In vitro* assays employing X-Tfi^XAC2610^ mutants were used to test and validate an AlphaFold2-derived model of the X-Tfe^XAC2609^-X-Tfi^XAC2610^ complex which presents topological similarities with the distantly related Tse1/Tsi1 complex from *P. aeruginosa* and the the i-type lysozyme from *Meretrix lusoria* (MI-iLys) in complex with PliI-Ah from *Aeromonas hydrophila*. While immunity proteins in other systems have been shown to protect against attacks by sister cells (*trans*-intoxication), this is the first description of an antibacterial secretion system in which the immunity proteins are dedicated to protecting cells against *cis*-intoxication.

## Introduction

Bacteria are continuously competing with each other for space and nutrients. One widespread competition strategy is the secretion of effectors into adjacent bacterial cells (Russell et al. 2011; Benz and Meinhart 2014; Souza et al. 2015; Cianfanelli, Monlezun, and Coulthurst 2016) mediated by specialized secretion systems, as for example: the Type IV Secretion Systems (T4SSs) from *Xanthomonas citri* (Souza et al. 2015) and *Stenotrophomonas maltophilia* (Bayer-Santos et al. 2019), the Type V Secretion System (T5SS) from *Escherichia coli* (Aoki et al. 2005), the Type VI Secretion Systems (T6SSs) from *Pseudomonas aeruginosa* (Russell et al. 2011), *Burkholderia thailandensis* (Schwarz et al. 2010), *Salmonella typhimurium (Sana et al. 2016)*, *Vibrio cholerae (MacIntyre et al. 2010)* and *Serratia marcescens* (Murdoch et al. 2011), the *Caulobacter crescentus* CDZ-based Type I secretion System (García-Bayona, Guo, and Laub 2017), and the Type VII Secretion Systems (T7SSs) encoded by Gram-positive bacteria *Staphylococcus aureus* (Cao et al. 2016) and *Bacillus subtilis* (Tassinari et al. 2022; Kobayashi 2021). All these different secretion systems have at least one effector and one cognate immunity protein, the latter of which protects the donor from self-intoxication.

T4SSs are large protein complexes which traverse the cell envelope, producing a channel through which proteins or protein–DNA complexes can be secreted into animal or plant hosts or other bacterial cells (Alvarez-Martinez and Christie 2009; Ilangovan, Connery, and Waksman 2015; Y. G. Li and Christie 2018; Sheedlo et al. 2022). Canonical type-A T4SSs are usually composed of 12 proteins, VirB1-VirB11 and VirD4 (Fronzes, Christie, and Waksman 2009; Alvarez-Martinez and Christie 2009; Costa et al. 2015). Chromosome-encoded T4SSs found in the order Xanthomonadales and some other proteobacterial species (X-T4SSs) secrete antibacterial effectors (X-Tfes) that are recruited via interactions with the VirD4 ATPase coupling protein (Alegria et al. 2005; Souza et al. 2011; Oka et al. 2022; Sgro et al. 2019). All X-Tfes contain a conserved C-terminal domain termed XVIPCD that interacts directly with the all-alpha domain of VirD4 (Alegria et al. 2005; Oka et al. 2022). In the phytopathogen *Xanthomonas citri*, the X-Tfe^XAC2609^ effector is secreted in a manner that is dependent on its XVIPCD and on a functional X-T4SS (Souza et al. 2015; Oka et al. 2022). The N-terminal portion of X-Tfe^XAC2609^ contains a glycoside hydrolase family 19 (GH19) domain that cleaves peptidoglycan (PG) and a PG binding domain. This PG hydrolase activity is inhibited by its cognate immunity protein X-Tfi^XAC2610^ (Souza et al. 2015). Therefore, X-Tfe^XAC2609^ and X-Tfi^XAC2610^ form an effector/immunity protein pair associated with the *X. citri* X-T4SS, which provides an adaptive advantage for X. *citri* in co-cultures with *E. coli* and other bacterial species (Oka et al. 2022; Souza et al. 2015).

Secretion system-mediated bacterial killing is typically evaluated in interbacterial competition assays between attacker and prey cells that code for one or more different effector-immunity protein pairs. The rationale is that the prey is susceptible to the toxicity of the delivered effector(s) because they do not produce at least one cognate immunity protein (Aoki et al. 2005; Russell et al. 2011, 2012; García-Bayona, Guo, and Laub 2017; Kobayashi 2021; Tassinari et al. 2022). Since most bacteria-bacteria interactions are between genetically identical cells (e.g. bacterial colonies), bacteria coding for toxic effectors also code for immunity proteins that protect themselves against intoxication via their own effectors. It is reasonable to suppose that immunity proteins should be localized in the same subcellular compartment where its cognate effector acts (Benz and Meinhart 2014; Whitney et al. 2013; Russell, Peterson, and Mougous 2014; Jurėnas and Journet 2021); for example, the X-Tfe^XAC2609^ toxin that targets peptidoglycan has a cognate immunity protein, X-Tfi^XAC2610^, that carries an N-terminal signal peptide and lipobox that directs it to the periplasm (Souza et al. 2015; Sgro et al. 2019). In this scenario, immunity proteins could, in principle, provide protection against two different, *not necessarily exclusive*, toxicity mechanisms: i) intoxication due to “friendly-fire” translocation of toxic effectors from neighboring sister cells (fratricide or *trans*-intoxication; **Figure 1A**) or ii) self-intoxication (suicide or *cis*-intoxication; **Figure 1B**) that results from the action of endogenously produced toxins. Both *cis-* and *trans*-intoxication mechanisms have been observed previously for T6SS mediated effector transfer (Basler and Mekalanos 2012; Hood et al. 2010; Russell et al. 2011; Dong et al. 2013); (M. Li et al. 2012; Whitney et al. 2015); (Basler and Mekalanos 2012; Hood et al. 2010; Russell et al. 2011; Dong et al. 2013).

**Figure 1.**
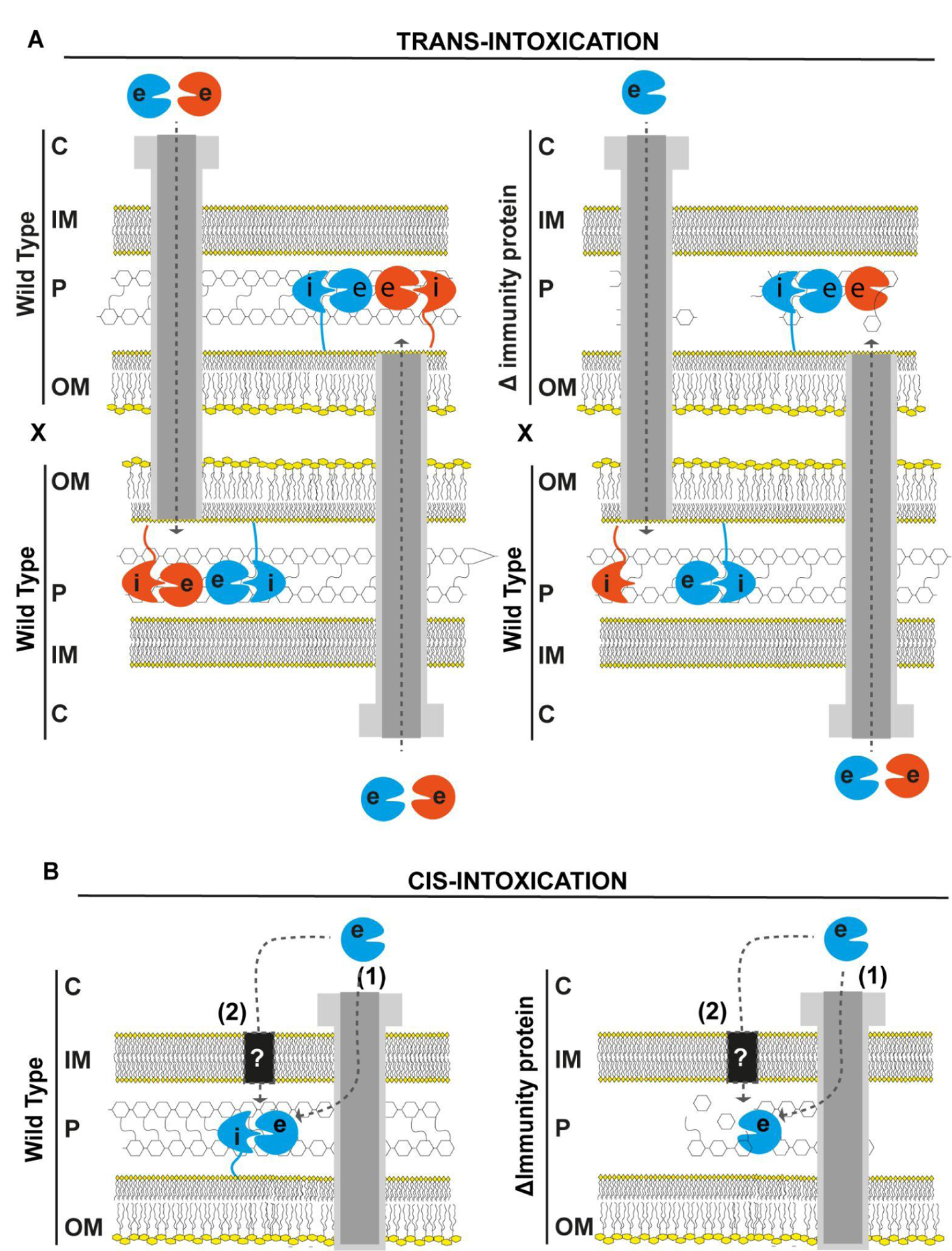
Schematic model of *trans-* and *cis-*intoxication mechanisms. **(A)** *Trans-*intoxication. In this mechanism, intoxication is due to contact- and T4SS-dependent transfer of X-Tfes (effectors) from one cell to another. *Left:* G enetically identical cells with equivalent repertoires of X-Tfes and cognate X-Tfis (immunity proteins) would be protected. In the scheme shown here, two wild-type cells that produce two different effector-cognate immunity protein pairs (orange and blue; e: effectors and i: immunity proteins) are immune against the toxic effects of the X-T4SS-mediated *trans*-intoxication due to the protective role of cognate immunity proteins. *Right:* Encounters between cells with non-equivalent repertoires would lead to killing. In the scheme shown here, a wild-type cell that produces two different effector-cognate immunity protein pairs (blue and orange) is in contact with a mutant cell that produces only one effector-immunity protein pair (blue). The hypothesis is that the two cells transfer effectors into each other’s periplasm and since the prey cell lacks the immunity protein that inhibits the orange effector, its cell wall is susceptible to degradation. **(B)** *Cis*-intoxication. In this mechanism, instead of being transported outside of the cell by the X-T4SS, an effector is translocated into the periplasm. Translocation could be T4SS-dependent (1) or T4SS-independent (2). *Left*: A wild type cell carrying a complete set of cognate immunity proteins is protected against self intoxication. *Right:* A bacterial strain lacking the immunity protein (ΔImmunity protein), may be susceptible to the cumulative activity of an effector that leaks into the periplasm. Cytoplasm (C); Inner membrane (IM); outer membrane (OM); Periplasm (P).

Here, we show that a *X. citri* strain lacking multiple effector-immunity protein pairs remains resistant to fratricidal X-T4SS-mediated attack by wild-type *X. citri* cells, thus providing evidence that the role of X-T4SS immunity proteins (X-Tfis) is not restricted to avoiding X-T4SS-mediated fratricide (*trans*-intoxication). We also show that a *X. citri* X-Tfi^XAC2610^ knockout strain suffers autolysis in a process that is mediated by X-Tfe^XAC2609^ but is independent of the X-T4SS. Cell ultra-structural aspects of the autolysis process were analyzed by fluorescence and electron microscopies. We demonstrate that X-Tfi^XAC2610^ is important for biofilm formation and is required to protect against the detrimental effects of X-Tfe^XAC2609^, even in the absence of a functional X-T4SS. These results support the conclusion that the protective function of X-Tfis is geared mainly towards the *cis*-intoxication (self-intoxication) effects of the endogenous X-Tfes.

## Materials and Methods

### Cultivation conditions

The oligonucleotides, plasmids and bacterial strains used in this work are described in Tables S1, S2 and S3, respectively. *X. citri* strains were cultivated in LB-agar or 2xTY media. The concentrations of the antibiotics used were: 70 μg/mL ampicillin, 100 μg/mL spectinomycin, 20 μg/mL gentamicin and 100 μg/mL kanamycin. Experiments involving *X. citri* were initiated by picking isolated colonies for inoculation in 5 mL of 2xTY medium supplemented with antibiotics and grown for 12 hours. The cells were then harvested, the optical density (OD_600nm_) adjusted to 0.05 with fresh 2xTY medium and a 2 mL volume of this culture was grown at 30°C, 200 rpm for another 12-18 hours.

### Production of *X. citri* gene-knockout strains

Based on the genomic DNA sequence of *X. citri* strain 306 (da Silva et al. 2002), oligonucleotides (Table S1) were designed to create the desired pNPTS-derived suicide vectors (Table S2). These vectors were used to produce the *X. citri* simple △X-Tfi^XAC2610^, △VirB4, △VirB5, △VirB6, △VirB8, and double △X-Tfi^XAC2610^/Vir(B(4-10), D4) and △X-Tfi^XAC2610^/△X-Tfe^XAC2609^ knockout strains using a two-step allelic exchange procedure, as previously described for the △VirB7, △VirB9, △VirB10 and △X-Tfe^XAC2609^ strains (Souza et al. 2015, 2011; Sgro et al. 2018). The mutants were confirmed by sequencing, or by colony PCR and Western Blot.

### Cloning, site-directed mutations and complementation

From the genomic sequence of *X. citri*, oligonucleotides were designed for complementation of the △X-Tfi^XAC2610^ strain (Table S1). The PCR products were cleaved with restriction enzymes NcoI and SalI, and inserted into the pBRA plasmid vector (M. Marroquin, unpublished). Site-directed mutation was performed using the QuikChange II XL kit (Agilent). Pairs of oligonucleotides F_XAC2609-E48A/R_XAC2609-E48A, F_2610-Y170A/R_2610-Y170A (Table S1) and plasmids pBRA-2609Nt, pET28a-XAC2610His-22-267 were used to construct the vectors pBRA-XAC2609NtE48A and XAC2610His-22-267, respectively (Table S2). *X. citri* wild-type and mutant strains were transformed by electroporation (2.0 kV, 25 μF, 200 Ω) and isolated colonies were selected on LB-agar medium containing 100 μg/mL spectinomycin, 70 μg/mL ampicillin.

### Western blotting

After the cultivation of the *X. citri* inocula in 2xTY supplemented with spectinomycin, cells were harvested by centrifugation (5,000 rpm, 5 minutes) and resuspended in water to an OD_600nm_ of 40. A 5-µL aliquot of resuspended cells was lysed in SDS-PAGE loading buffer at 100°C for 5 minutes, separated by SDS-PAGE and assayed by Western blot. Rabbit polyclonal antibodies serum at 1:1,000 dilution anti-X-Tfe^XAC2609^, anti-X-Tfi^XAC2610^ and anti-VirB7 (Souza et al. 2015) were detected by IRDye 800 CW goat anti-rabbit IgG (LI-COR Biosciences) at 1:30,000 dilution. Secondary antibody signals were detected using an Odyssey infrared imaging system (LI-COR Biosciences).

### Colony transparency assay

After the cultivation of the *X. citri* inocula in 2xTY medium supplemented with spectinomycin, cells were harvested by centrifugation (5,000 rpm, 5 minutes), washed in water twice and the OD_600nm_ was normalized to 0.05 in 2xTY medium. Then, 5 µL of each *X. citri* strain culture was applied onto the surface of 1.5% LB-agar plates supplemented with 70 μg/mL ampicillin, spectinomycin and 0.1% arabinose. The plates were cultured at 30°C and photographs were recorded on a transilluminator every 24 hours.

### Colony viability assay

After the cultivation of the *X. citri* inocula in 2xTY supplemented with spectinomycin and 0.1 % arabinose, cells were harvested by centrifugation (5,000 rpm, 5 minutes), washed in water twice and the OD_600nm_ was normalized to 0.05. Then, 5 µL of each *X. citri* strains were pipetted onto 1.5 % LB-agar plates supplemented with spectinomycin and 0.1% arabinose. Next, the plates were cultured at 30°C. To assess the cell viability, *X. citri* colonies were removed from the LB-agar plates and gently resuspended in 1 ml of 2xTY, every 24 hours of cultivation. Serial dilutions were made on LB-agar (1.5%) plates supplemented with ampicillin and spectinomycin to estimate viability of colony forming units per ml (CFU/mL). The results shown are the means of 3 independent experiments.

### Transmission electron microscopy (TEM)

After growth, liquid cultures of *X. citri* cells were centrifuged in microtubes (5,000 rpm, 5 minutes, room temperature) and supernatants were discarded. Cell washes were performed twice by adding 1 mL of phosphate buffer solution (0.2 M, pH 7.4). After 15 minutes at room temperature, samples were centrifuged (6,500 rpm, 5 minutes) and the supernatant discarded. A fixation step was performed by the addition of 1 mL of modified Karnovsky’s solution (2.5% glutaraldehyde, 2% paraformaldehyde, 200 mM phosphate buffer, pH 7.4) (Watanabe and Yamada 1983). Then, the samples were incubated for 90 minutes at room temperature and centrifuged (5,000 rpm, 5 minutes, room temperature), the supernatant was discarded and an extra washing step was carried out with phosphate buffer (200 mM, pH 7.4). A second fixation step was performed by the addition of 1 mL of 1% OsO_4_ solution followed by incubation for 1 hour on ice. The samples were then centrifuged (5,000 rpm, 2 minutes, room temperature) and washed with water. The pellets were dissociated from the microtubes with the use of a glass rod. Thereafter, 1 mL of 0.5% aqueous solution of uranyl acetate was added, followed by another incubation period of 1 hour at room temperature.Next, uranyl acetate was removed by centrifugation and 1 mL of 60% ethanol solution was added. Successive incubations were performed with increasing concentrations of ethanol (70%, 80%, 90%, 95% and 100%). After the last dehydration step with 100% ethanol, 1 mL of propylene oxide solution (Electron Microscopy Sciences) was added, followed by incubation at room temperature for 10 minutes. Cells were collected by centrifugation as above and three more incubations and washes with propylene oxide were performed. Then, the samples were submitted to 4 successive exchanges of propylene oxide and resin solutions for microscopy (Low Viscosity embedding Kit (# 14300), Spurr’s) in proportions of 1:1, 1:3, 0:1 and 0:1 (propylene oxide:resin) with incubation times of 40 minutes, 90 minutes and 12 hours and 72 hours, respectively. The first 3 incubations (1:1, 1:3 and 0:1) were performed with gentle shaking at room temperature. The fourth incubation step was performed at 60°C without agitation in order to solidify the resin. After resin solidification, histological and ultrathin sections were obtained. The slices were collected on 200 mesh carbon-covered copper grids (Ted Pella). Sample imaging was performed on JEM 2100 JEOL (Central Analitica, Institute of Chemistry, University of São Paulo) and JEOL 100 CX II (Institute of Biomedical Sciences, University of São Paulo) transmission microscopes. Quantitative analysis of the number of cells having a damaged cell envelope and cell counting were performed by manual inspection of the micrography images.

### Time-lapse fluorescence microscopy

Time-lapse fluorescence microscopy assays were carried out as previously described (Oka et al. 2022). Briefly, after the cultivation of the *X. citri* inocula in 2xTY supplemented with appropriate antibiotic, cells were harvested by centrifugation (5,000 rpm, 5 minutes), washed in water twice and the OD_600nm_ was normalized to 0.5. Then, 1 µL of each *X. citri* strain were pipetted onto a thin LB-agarose support supplemented with propidium iodide (1 µg/mL), appropriate antibiotic and observed with a Nikon Eclipse Ti microscope equipped with filters for GFP (GFP-3035B-000-ZERO, Semrock) and propidium iodide (TxRed-4040B, Semrock) and a Nikon Plan APO 100x objective. Images were collected every 10 minutes. Image processing and quantitative analysis of the number of cells having a damaged cell envelope and cell counting related to Smovies (1-5) were performed manually using Fiji software (Schindelin et al. 2012) multipoint tool. Video microscopy showing interbacterial competition between *X. citri* strains wild type transformed with pBBR-RFP against *Δ8Δ2609-GFP transformed with pBBR-GFP,* the after cells growing and washing steps, the cultures were mixed 1:1 and the microscopy was performed as described above, except by supplementing the LB-agarose support with propidium iodide.

### Convolutional Neural Network (CNN) analysis

A pre-trained MobileNet CNN (Howard et al. 2017) was trained using a dataset containing 50 distinct images of both opaque and transparent colonies (with multiple instances per photo). The model was validated with 23 images of opaque colonies and 26 images of transparent colonies from our colonies. Confidence index for predictions was computed using the np.argmax(prediction) function from the NumPy library (version 1.23.5). Confidence level associated with each prediction was accessed using prediction[0][class_idx]. Tensor Flow optimization, data augmentation, and learning rate reduction were employed for training the CNNs.

### Protein expression and purification

X-Tfi^XAC2610^His-22-267 expression and purification were carried out as previously described (Oka et al. 2022), with some adaptations. Briefly, cells of *E. coli* Bl21 (DE3) carrying the vectors (Table S2) were grown to OD_600nm_ = 0.8 and the heterologous expression of recombinant proteins was induced using 0.5 mM IPTG for 16 hours at 18°C and 180 rpm in 2xTY medium. After induction of protein expression, cell recovery and lysis, the proteins of interest were submitted to affinity chromatography using HiTrap Ni^2+^-chelating resin (Cytiva) previously equilibrated with 20 mM Tris-HCl buffer (pH 8.0), 200 mM NaCl, 20 mM imidazole, and 2% (v/v) glycerol, subsequently washed with the same buffer and eluted with an imidazole gradient (20-500 mM). X-Tfe^XAC2609^(1-308) was purified by chromatography as previously described (Souza et al. 2015), using an anion exchange Q-sepharose column (GE Healthcare) and a size exclusion Superdex S200 column (GE Healthcare). Pure proteins were identified by SDS-PAGE, and the quantification was estimated using absorbance at 280 nm.

### *Micrococcus luteus* peptidoglycan hydrolysis assay

Peptidoglycan hydrolysis experiments were performed as previously described (Souza et al. 2015), with some modifications. Briefly, *M. luteus* cell wall (Sigma) suspensions at OD_600nm_ of 0.7 in 50 mM sodium acetate (pH 5.0) and 2 mM CaCl_2_ were incubated in triplicate for 60 minutes at 30°C with only buffer (negative control), 2 μM X-Tfe^XAC2609^(1-308), 2 μM X-Tfe^XAC2609^(1-308) plus 2 μM X-Tfi^XAC2610^(His-55-267), and 2 μM X-Tfe^XAC2609^(1-308) plus 2 μM of X-Tfi^XAC2610^(His-55–267) Y170A. After 10, 20, and 60 minutes of incubation, the reactions were quenched by adding 500 mM sodium carbonate. The decrease in optical density due to peptidoglycan hydrolysis was monitored over time at 600 nm.

### Biofilm formation assay

*X. citri* inocula containing pBRA- and pBBR-derived plasmids were grown for 12 hours in 2xTY supplemented with ampicillin, gentamicin, spectinomycin and 0.3% arabinose. After growth, *X. citri* inocula OD_600nm_ was normalized to 0.05 and incubated for 5 days at 30°C in a laminated microscopy chamber (Nu155411; Lab-Tek, NUNC). Biofilm images were acquired using a Nikon Eclipse Ti microscope equipped with a 100x magnification objective (CFI Plan Apo Lambda 100XH) and a fluorescence filter for GFP (GFP-3035B, Semrock). Images were collected from the base to the top over 20 µm in the Z axis at 0.5-µm intervals, and stacked using the FIJI software (Schindelin et al. 2012). The results shown are representative images of 3 independent experiments. For the 24 wells plate-based biofilm assay, *X. citri* inocula were grown for 24 hours at 30°C, 200 rpm in 2xTY medium supplemented with ampicillin, and then allowed to grow at room temperature (22°C) for seven days without shaking. To quantify *X. citri* biofilm using a 24 well plate, the protocol described in (Dunger et al. 2014) with some modifications was followed. After seven days of cultivation, the 2xTY medium was carefully removed and replaced with 1 mL of 0.1M NaCl solution. The cells were then resuspended by pipetting and transferred into 1.5 mL microtubes. After vortexing and centrifuging (6000 rpm, 5 minutes at 4°C) the supernatant was discarded, this step was repeated two more times. Next, 1 mL of 0.1% crystal violet solution was added, mixed by vortexing, and incubated for 30 minutes at room temperature. The tubes were then centrifuged (6000 rpm, 5 minutes at 4°C), the supernatant was discarded, and the cells were washed two more times with 1 mL of 0.1M NaCl followed by centrifugation steps. Finally, the pellets were resuspended with 100% ethanol and the absorbances were measured at 570 nm.

### Fluorescence spectroscopy

The protein X-Tfi^XAC2610^(55-267) was expressed and purified as previously described (Souza et al. 2015). Fluorescence assays of the purified protein were measured employing an ATF-105 spectrofluorometer (Aviv Biomedical). Thermal denaturation experiments were conducted with X-Tfi^XAC2610^(55-267) at 0.5 mM in 20 mM Tris-HCl (pH 7.5) and 50 mM NaCl. Where indicated, EGTA, salts (MgCl_2_ and CaCl_2_) were added with final concentrations of 0.5 mM and 0.75 mM, respectively. The temperature was increased between 20°C and 90°C, with steps of 2°C and 6 minutes equilibration per step. Intrinsic tryptophan fluorescence was excited at 295 nm (bandwidth of 2 nm) and emission was detected at 337 nm (bandwidth of 5 nm). The fraction of folded protein was calculated as described in (Pace and Scholtz 1997)

### Bioinformatics

Prediction of the structure of the X-Tfi^XAC2610^(54-267)/X-Tfe^XAC2609^(1-194) complex was performed using the UCSF ChimeraX (v1.4, 2022-06-03) software that includes an integrated link to the ColabFold-AlphaFold2 suite (Mirdita et al. 2022; Jumper et al. 2021; Varadi et al. 2022). The search for homologous proteins was performed using the Blast algorithm (Dooley 2004) with X-Tfi^XAC2610^(54-267) as a query against refseq NCBI protein data bank using an expect values threshold of 1×10^-7^, and outliers and sequences with 99% redundancy were manually discarded. Next, the sequences were realigned with Muscle (Edgar 2004) using the X-Tfi^XAC2610^ sequence as a reference. The realigned sequence file (S1 Supplementary File) was used as input for the Weblogo software (Crooks et al. 2004) to create the conservation profile. Structures were rendered using UCSF Chimera (v1.16) and ChimeraX (v1.4)(Pettersen et al. 2021).

### Citrus canker assay

Citrus canker assays were performed as previously described (Souza et al. 2011). Briefly, *X. citri* inocula were grown for 12 hours in 2xTY supplemented with ampicillin. After growth, *X. citri* cells diluted in PBS buffer (137 mM NaCl, 2.7 mM KCl, 8 mM Na_2_HPO_4_, 2 mM KH_2_PO_4_) to an OD_600nm_of 0.1, and 100 µl used to infiltrate sweet orange leaves (*Citrus sinensis* (L.) Osbeck) using a syringe with a needle. Plants were maintained at 28°C with a 12 hour photoperiod, and symptom development was regularly observed and recorded.

### *X. citri* vs *E. coli* competition assays

Chlorophenol-red β-D-galactopyranoside (CPRG)-based bacterial competition assays were performed as previously described (Oka et al. 2022). Briefly, *X. citri* at OD_600nm_of 2.0 were mixed 1:1 with *E. coli* MG1655 at OD_600nm_ of 11. The mixed cultures (5 μL) were spotted on 1.5% agarose supplemented with 40 µg/mL CPRG (Sigma-Aldrich) in 96 wells plates. Absorbance at 572 nm was monitored at each 10 minutes using a plate reader (SpectraMax Paradigm, Molecular Devices).

### *X. citri* vs *X. citri* competition assays

For the bacterial competition assay based on viability, *X. citri* harboring pBBR5GFP or pBBR2RFP were grown at 30°C for 12 hours in 2xTY supplemented with gentamicin (20 µg/mL) or kanamycin (50 µg/mL), respectively. After three washing steps with fresh 2xTY, the *X. citri* cultures at OD_600nm_ of 2.0 were mixed 1:1, and five microliters were spotted in 1% LB-agar plates supplemented with ampicillin (100 µg/mL) for 40 hours at 30°C. Finally, the colonies were retrieved and resuspended in 2xTY medium, and the cellular viability per colony was estimated by serial dilution using selective medium supplemented with kanamycin or gentamicin in LB agar plates.

## Results

### The X-T4SS immunity proteins are not the primary defense against *trans*-intoxication (fratricide)

Figure S1 shows that *X. citri* is able to kill *E. coli* in an X-T4SS dependent manner, as has been previously shown (Souza et al. 2015; Oliveira et al. 2016; Oka et al. 2022). Figure 2A shows CPRG-based colorimetric assays to quantitatively monitor real-time killing of *E. coli* cells in *X. citri/E. coli* co-cultures. As previously shown, the ΔVirB7 and *X. citri Δ8Δ2609-GFP* strains do not kill *E. coli (Bayer-Santos et al. 2019; Oka et al. 2022)* demonstrating that the X-T4SS presents antibacterial activity and that this activity is dependent on the presence of a cohort of secreted effectors (X-Tfes). On the other hand, the deletion of X-Tfi^XAC2610^ or the X-Tfe^XAC2609^/X-Tfi^XAC2610^ pair does not significantly impair the antibacterial function of the X-T4SS under the conditions tested. In order to test whether the *trans*-intoxication (fratricide) hypothesis (**Fig 1A**) is valid for the X-T4SS, we performed intraspecies bacterial competition assays using wild-type *X. citri* against its derivative mutants. **Fig 2B** shows that the X-T4SS fails to confer a competitive advantage against the *X. citri* Δ*virB7* strain that does not produce a functional X-T4SS due to the absence of the VirB7 subunit (Souza et al. 2015; Oliveira et al. 2016; Sgro et al. 2018). This result itself is not inconsistent with the *trans*-intoxication hypothesis since the target *X. citri* Δ*virB7* strain still carries a full set of X-Tfis. However, wild-type *X. citri* cells were also unable to kill the ΔX-Tfe^XAC2609^/ΔX-Tfi^XAC2610^ double-mutant *X. citri* strain which lacks the X-Tfe^XAC2609^-X-Tfi^XAC2610^ toxin-antitoxin pair (Figure 2B) nor do they kill the *Δ8Δ2609-GFP X. citri* strain in which eight other X-Tfe/X-Tfi effector/immunity protein pairs were deleted (XAC2885/XAC2884, XAC0574/XAC0573, XAC0096/XAC0097, XAC3634/XAC3633, XAC1918/XAC1917, XAC0466/XAC0467, XAC4264/XAC4263/XAC4262, XAC3266/XAC3267; (Oka et al. 2022)). The observation that wild-type *X. citri* is unable to kill the ΔX-Tfe^XAC2609^ΔX-Tfi^XAC2610^ or *X. citri Δ8Δ2609-GFP* strains indicate that the primary role of X-T4SS immunity proteins is not to neutralize exogenous effectors that were injected by neighboring bacteria (*trans*-intoxication) as exemplified in **Fig 1A**.

**Figure 2.**
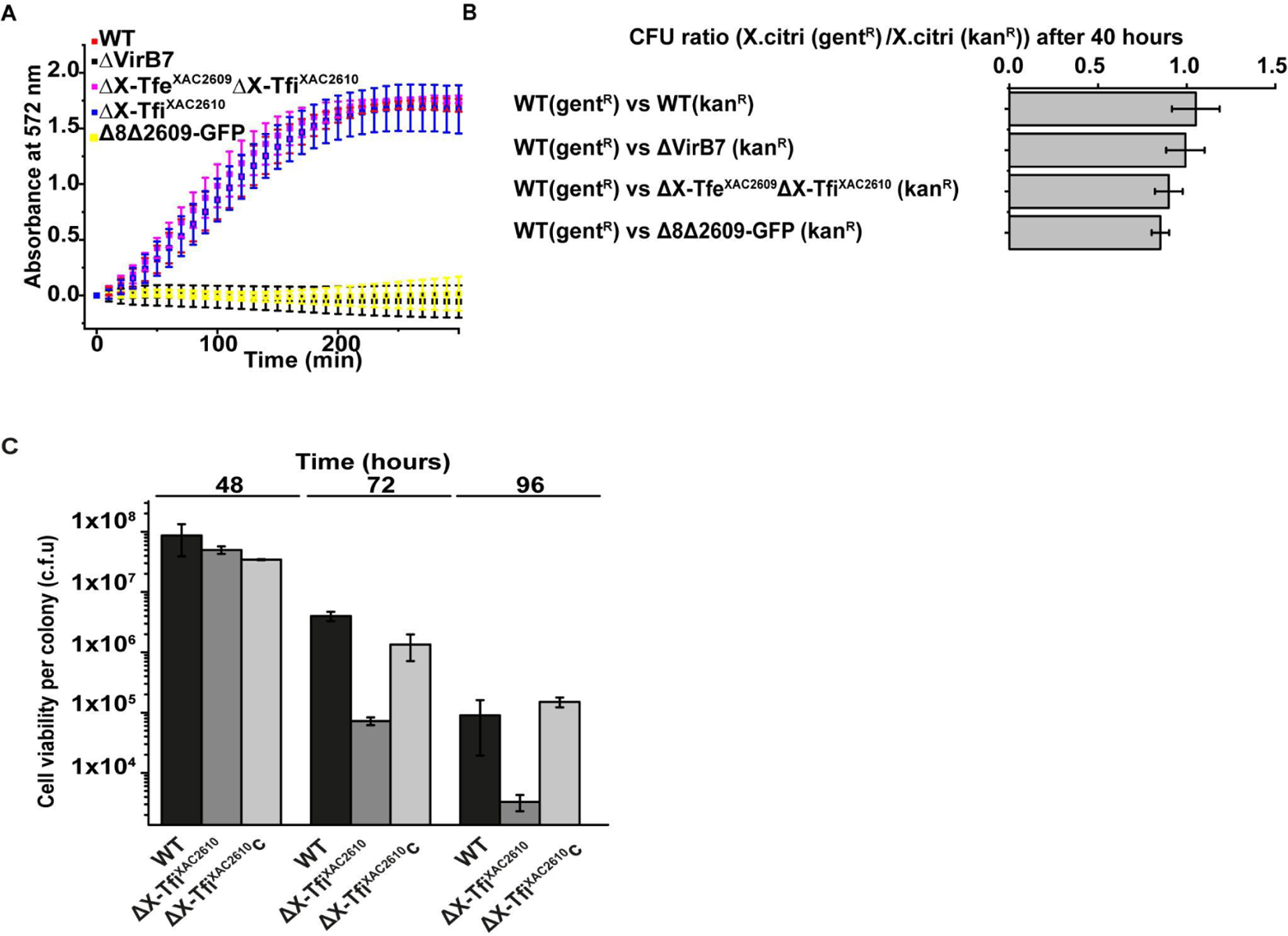
X-T4SS-related immunity proteins are not required to confer protection against *trans-*intoxication mediated by the X-T4SS. (A) Bacterial competition of *X. citri* against *E. coli* MG1655 cells expressing β-galactosidase. *E. coli* killing was monitored by detecting the degradation of chlorophenol red-β-D-galactopyranoside (CPRG) by measuring absorbance at 572 nm at 10-min intervals. (B) *X. citri* cell viability ratio after co-culture in Luria-Bertani (LB) agar after 40 hours. *X. citri* strains used: wild type (WT), ΔvirB7 (ΔVirB7), ΔX-Tfe^XAC2609^ΔX-Tfi^XAC2610^, and the *Δ8Δ2609-GFP* carrying the vectors pBBR(MCS5) GFP and pBBR(MCS2) RFP and that confer resistance to gentamicin (gent^R^) and kanamycin (kan^R^). Colony forming units per mL (CFU/mL) of each strain was assessed through serial dilution assays on LB agar plates carrying the appropriate antibiotics. Error bars represent the SD of three experiments (one way ANOVA p-value = 0.291).**(D)** Colony viability assay. *X. citri* wild-type (WT) strain (black), ΔX-Tfi^XAC2610^ (dark gray) strain, and ΔX-Tfi^XAC2610^ + X-Tfi^XAC2610^ strain (gray) strain were grown on LB agar plates. After 48 hours,72 hours, and 96 hours at 30 °C, the colonies were resuspended in 2xTY media, and cellular viability (CFU/mL) was assessed through serial dilution assays on LB agar plates. Errors bars, s.d.; n= 3.

### X-Tfi^XAC2610^ provides immunity against *in vivo* intracellular autolytic activity of X-Tfe^XAC2609^

As the above results are inconsistent with the proposition that X-Tfis provide protection against *trans*-intoxication, we performed experiments to test the hypothesis that they instead provide protection against self-intoxication (*cis*-intoxication), as illustrated in **Fig 1B**. In the case of effectors that normally act in the periplasm of target cells (for example, lysozyme-like effectors), *cis*-intoxication may arise if the effector is transferred across the inner membrane and into the periplasm of the producing cell by one or more routes, either dependent or independent of the X-T4SS. To test this hypothesis, we created *X. citri* strains with single or multiple in-frame deletions of genes coding for the X-Tfe^XAC2609^ lysozyme-like effector, its cognate immunity protein X-Tfi^XAC2610^ and the VirB7 and VirD4 subunits that are essential for X-T4SS function.

Figure S2 show that colonies of *X. citri* wild-type, ΔX-Tfi^XAC2610^, ΔX-Tfi^XAC2610^Δ*virB7*, ΔX-Tfi^XAC2610^c (c: complementation with an extrachromosomal plasmid expressing X-Tfi^XAC2610^), and ΔX-Tfi^XAC2610^cΔ*virB7* grown on LB-agar plates for 24 and 48 hours are indistinguishable in terms of color, roughness, opacity and size. However, after 72 hours of growth, colonies of all strains with X-Tfe^XAC2609^ but lacking X-Tfi^XAC2610^ (ΔX-Tfi^XAC2610^, ΔX-Tfi^XAC2610^Δ*virB7*) became partially transparent, indicative of cell death. On the other hand, the lineages lacking an X-Tfe^XAC2609^ gene or carrying an X-Tfi^XAC2610^ gene all maintained their opacity at 72 hours (including strains ΔX-Tfi^XAC2610^c, ΔX-Tfi^XAC2610^cΔ*virB7* that carry a plasmid that confers expression of X-Tfi^XAC2610^ and ΔX-Tfi^XAC2610^ transformed with pBRA-X-Tfi^XAC2610^His-22-267 that produces a cytoplasmic version of X-Tfi^XAC2610^. Importantly, no reduction in colony opacity was observed after 72 hours of growth for the double mutant ΔX-Tfe^XAC2609^ΔX-Tfi^XAC2610^, indicating that X-Tfe^XAC2609^ must be present for the development of the observed phenotype (**Fig. S2A**). Also of interest, the fact that a cytossolic version of X-Tfi^XAC2610^His-22-267 protects against X-Tfe^XAC2609^ toxicity suggest that complex formation in the cytoplasm may impede leakage into the periplasm. Finally, transparent colonies are observed even in the absence of VirB7 (Δ*virB7*), demonstrating that this phenotype does not depend on a functional X-T4SS, but rather results from the absence of the immunity gene. The colonies phenotypes shown were further validated using a convolutional neural network (CNN) analysis (**Fig S2**).

All genes studied in this manuscript are located in a single genomic locus, and knock-outs could potentially interfere with protein expression levels of nearby genes. To discard this possibility, Western blot assays were performed. These experiments showed that only the expression levels of the targeted genes are affected by the genetic manipulations **(Fig. S2B)**.

Figure 2C shows that after 48 hours of growth on LB agar, no difference in cell counting or viability between wild-type and ΔX-Tfi^XAC2610^ strains was observed. However, after 72 and 96 hours of growth, the ΔX-Tfi^XAC2610^ strain shows lower viability compared to the wild-type strain. The viability of the ΔX-Tfi^XAC2610^ strain was restored when complemented with an extrachromosomal plasmid expressing X-Tfi^XAC2610^.

In conclusion, these results indicate that i) the transparent colony phenotype is due to the activity of X-Tfe^XAC2609^, ii) it can be inhibited by the presence of X-Tfi^XAC2610^ and iii) the phenotype does not require a functional X-T4SS. These data are consistent with a *cis*-intoxication mechanism by X-Tfe^XAC2609^ that is inhibited by X-Tfi^XAC2610^.

### X-Tfi^XAC2610^ is important for maintaining the integrity of the *X. citri* cell envelope

The transparent colony phenotype described above suggests that, in the absence of its cognate immunity protein, the X-Tfe^XAC2609^ lysozyme-like effector induces cell autolysis. To test this hypothesis, we performed a set of time-lapse microscopy assays (Fig 3 and **Movies S1-5**) using wild-type and different *X. citri* mutant strains growing in media supplemented with propidium iodide (PI), a fluorescent dye that only stains nucleic acids when cell envelope integrity is compromised. We observed a significant increase in PI permeability in ΔX-Tfi^XAC2610^ cells (Fig 3**, Movie S2, Table S4**) compared to wild-type cells (Fig 3**, Movie S1. Table S4**). Permeability is reduced to wild-type levels in the ΔX-Tfe^XAC2609^ΔX-Tfi^XAC2610^ double mutant strain (Fig 3**, Movie S3, Table S4**). Transformation of ΔX-Tfe^XAC2609^ΔX-Tfi^XAC2610^ with a plasmid over-expressing the catalytic N-terminal domain of X-Tfe^XAC2609^(1-306), which lacks the XVIPCD X-T4SS secretion signal, greatly increased the amount of PI-permeable cells (Fig 3**, Movie S4, Table S4**). Moreover, multiple cell autolysis events were observed for the ΔX-Tfi^XAC2610^Δ*virD4 X. citri* strain (Fig 3**, Movie S5, Table S4**), confirming that toxicity caused by X-Tfe^XAC2609^ does not require the XVIPCD secretion signal and is not mediated by the X-T4SS. Close inspection of **Movies S2, S4, S5** and **S6** and Figure 4A shows that PI-permeability of *X. citri* cells coincides with a rapid change in cell morphology, from natural rod to spherical, consistent with X-Tfe^XAC2609^-induced weakening of the cell wall. The onset of PI permeability was observed to occur both in isolated cells and in cells in contact with neighbors (**Movie S6 and** Figure 4A). Another interesting observation is that the onset of PI-permeability was frequently observed in cells that were undergoing cell-division (**Movie S6**), perhaps coinciding with a phase in the cell cycle where peptidoglycan integrity is more susceptible to the deleterious hydrolytic activity of X-Tfe^XAC2609^.

**Figure 3.**
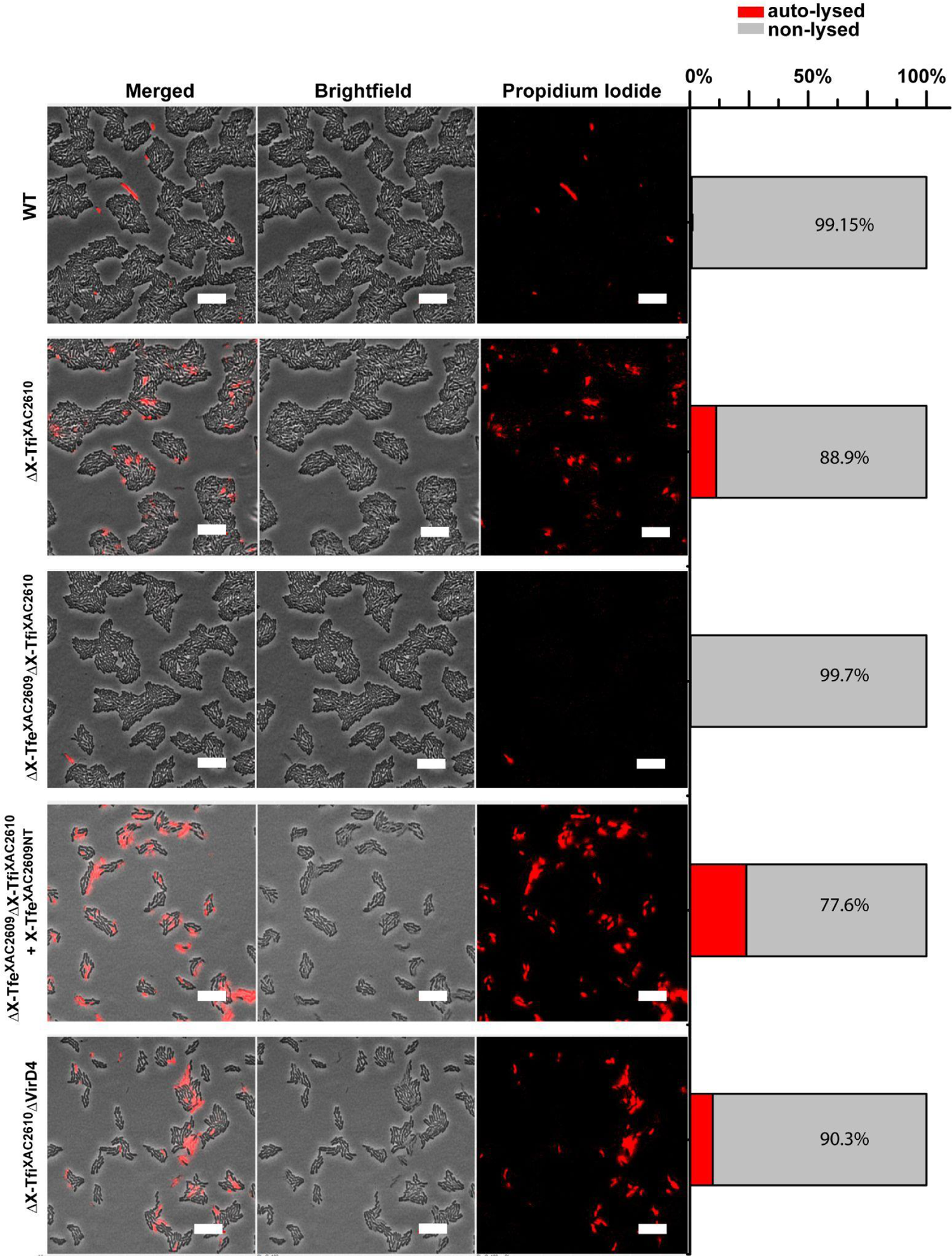
Quantitative analysis *of X. citri* propidium iodide permeability shown in Movies S1-S5. Micrographs show the last time point of each movie S(A)(1-5). Further details regarding quantitative analysis are presented in **Table S4**. Scale bar 10 µm.

**Figure 4.**
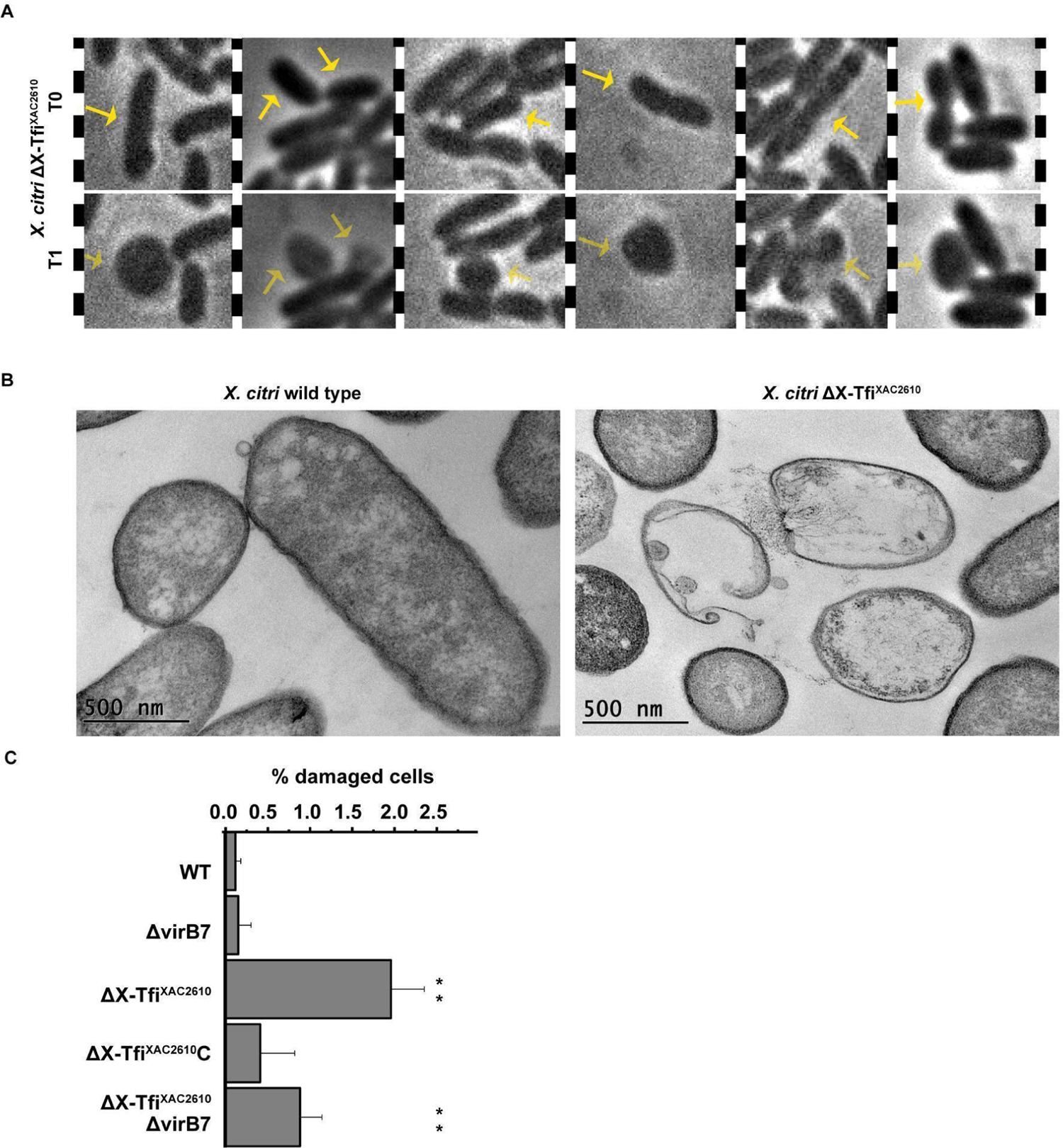
X-Tfi^XAC2610^ is important for maintaining *X. citri* cell envelope integrity. **(A)** *X. citri* ΔX-Tfi^XAC2610^ strain time-lapse microscopy shows spontaneous spheroplasts forming events. Arrows at T0 point to cells that will turn into spheroplasts at T1. See **movie S6** for more examples. **(B)** Transmission electron microscopy (TEM) micrographs of *X. citri* WT and ΔX-Tfi^XAC2610^ strains after 12 hours of growth in liquid 2xTY medium. Scale bars are 500 nm. (**C**) Percentage of damaged cells observed in TEM micrographs. More examples of damaged cells are shown in **Supplementary** Figures 3 and 4. *X. citri* strain used for the assay are described in the figure legends. *X. citri* strains used in the assay wild-type (WT) strain, ΔVirB7 strain, ΔX-Tfi^XAC2610^ strain, ΔX-Tfi^XAC2610^ΔVirB7, ΔX-Tfi^XAC2610^ strain complemented with X-Tfi^XAC2610^ (ΔX-Tfi^XAC2610^c). Means with a double asterisk are significantly different at P < 0.05 compared with the mean of the wild type, Tukey *post hoc* one-factor ANOVA test. Further details regarding quantitative and statistical analysis are shown in **Table S5.**

We then used transmission electron microscopy (TEM) in order to obtain a more detailed picture of the differences in the cell envelope ultrastructure of *X. citri* wild-type and ΔX-Tfi^XAC2610^ cells (Fig. 4B). TEM analyses of thin sections of previously fixed *X. citri* cells embedded in resin were used to assess structural details of the plasma membrane, cell wall and intracellular content. TEM micrographs of *X citri* ΔX-Tfi^XAC2610^ cells showed that they were frequently broken open with leakage of filamentous materials or were devoid of cellular contents (Fig. 4B and **S3, S4**). In contrast, wild-type cells typically present an intact cell envelope and a high-density intracellular environment (Fig. 4B and **Fig. S3, S4**). Micrographs of the ΔX-Tfi^XAC2610^ΔVirB7 strain presented a phenotype similar to that observed for ΔX-Tfi^XAC2610^ while micrographs of ΔVirB7 and ΔX-Tfi^XAC2610^c lineages showed mostly intact cells as observed for the wild-type strain (Fig. 4B and **Fig. S4**). Analysis of the number of intact versus damaged cells in TEM micrographs indicates significant statistical differences in the percentage of lysed cells in strains producing both X-Tfe^XAC2609^ and X-Tfi^XAC2610^ (wild-type, ΔVirB7 and ΔX-Tfi^XAC2610^c) compared with cells expressing the former but not the latter (ΔX-Tfi^XAC2610^ and ΔX-Tfi^XAC2610^ΔVirB; Fig. 4C**, S5 Table**). Together, fluorescence microscopy and TEM analyses indicate that, in the absence of X-Tfi^XAC2610^, the activity of X-Tfe^XAC2609^ promotes damage of the *X. citri* cell envelope, regardless of the presence of a functional X-T4SS.

### The hydrolytic activity of X-Tfe^XAC2609^ inhibits *X. citri* biofilm formation in the absence of X-Tfi^XAC2610^

The results so far show that the absence of X-Tfi^XAC2610^ compromises the integrity of the *X. citri* cell envelope of only a small number of actively dividing cells during the exponential growth phase (Fig. 3) and eventually leads to the lysis of a significant fraction of the cell population during stationary phase **(Fig S2)**. Since bacterial biofilms are a complex array of cells and extracellular polymers that develop over extended periods of time (many hours to days) with a progressive reduction in the rate of cell division (Dunger et al. 2014, 2016; Sena-Vélez et al. 2016), we decided to investigate the possible physiological effects associated with the deletion of X-Tfi^XAC2610^ on biofilm formation. Figure 5A shows that, unlike the *X. citri* wild-type strain, ΔX-Tfi^XAC2610^ and ΔX-Tfi^XAC2610^Δ*virB7* cells could not form biofilms on polystyrene plastic surfaces after five days of cultivation in 2xTY medium. The double knock-out strain ΔX-Tfe^XAC2609^ΔX-Tfi^XAC2610^ strain presents a normal biofilm while this double mutant strain complemented with either full-length X-Tfe^XAC2609^ or an N-terminal fragment lacking the XVIPCD secretion signal was defective in biofilm formation. On the other hand, the ΔX-Tfe^XAC2609^ΔX-Tfi^XAC2610^ strain complemented with the N-terminal domain of X-Tfe^XAC2609^ carrying a detrimental mutation (E48A) in the active site of its GH19 PG hydrolase domain (Souza et al. 2015) was able to form a normal biofilm. These results confirm that, in the absence of its cognate inhibitory protein, the deleterious effects of X-Tfe^XAC2609^ is dependent on its glycohydrolase activity.

**Figure 5.**
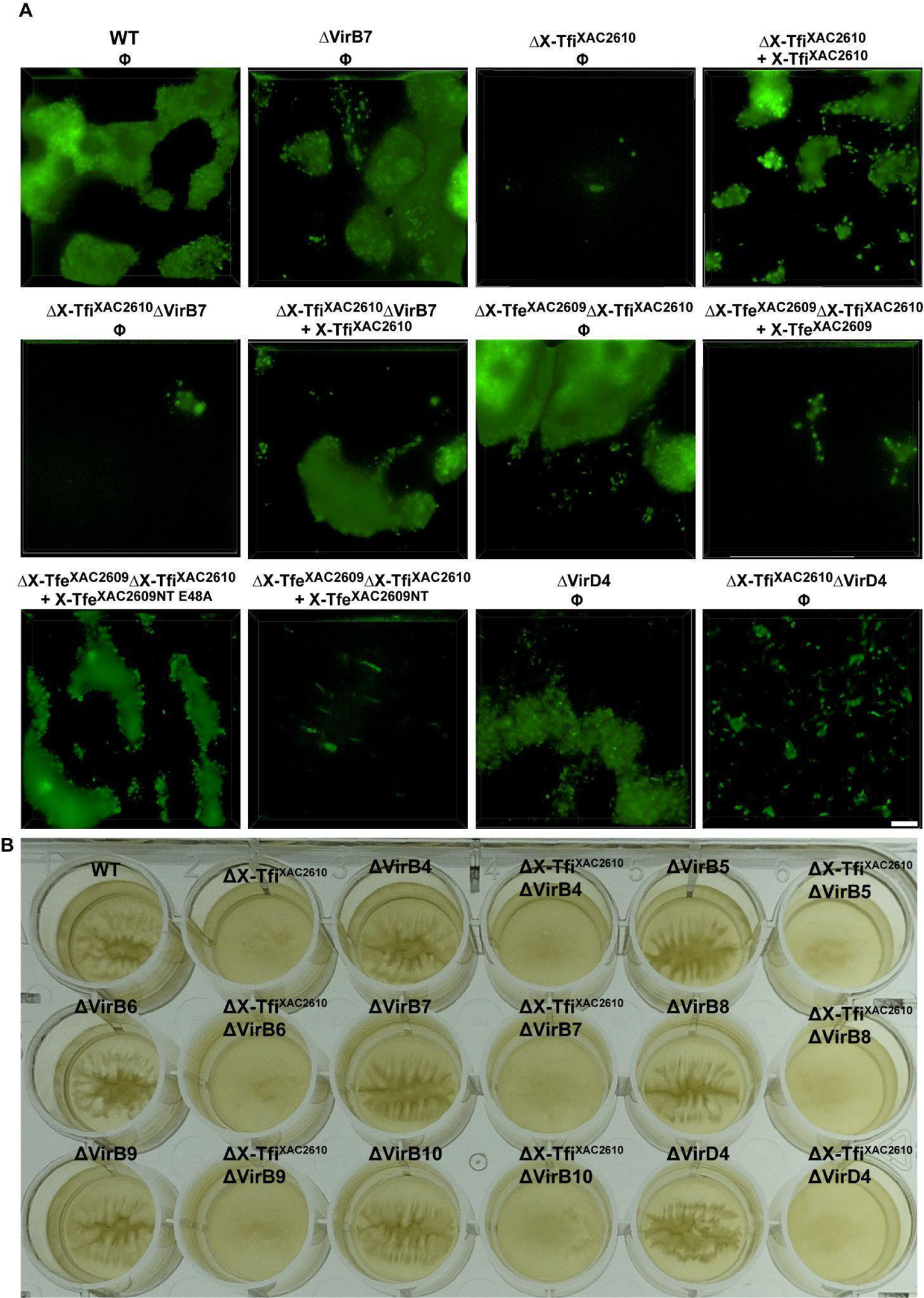
The hydrolytic activity of X-Tfe^XAC2609^ inhibits *X. citri* biofilm formation in the absence of X-Tfi^XAC2610.^ **(A)** *X. citri* wild type and derived mutants carrying a plasmid for the endogenous expression of GFP were grown on 2xTY medium for five days at 30 °C in chambered microscope slides. *X. citri* wild-type (WT) strain, ΔVirB7 strain, ΔX-Tfi^XAC2610^ strain, ΔX-Tfi^XAC2610^ΔVirB7 strain, ΔX-Tfe^XAC2609^ΔX-Tfi^XAC2610^ strain containing the empty vectors pBRA (Φ) and complemented strains (ΔX-Tfi^XAC2610^ + X-Tfi^XAC2610^, ΔX-Tfi^XAC2610^ΔVirB7 + X-Tfi^XAC2610^, ΔX-Tfe^XAC2609^ΔX-Tfi^XAC2610^ + X-Tfe^XAC2609^, _ΔX-Tfe_XAC2609_ΔX-Tfi_XAC2610 _+ X-Tfe_XAC2609NT_, ΔX-Tfe_XAC2609_ΔX-Tfi_XAC2610 _ΔX-Tfe_XAC2609_NT_ E48A) are indicated above each fluorescence microscopy image. Images were taken with a fluorescence microscope at 100X magnification. Scale bars, 5 µm. **(B)** Biofilm formation assay in 24 wells plates. Plates containing start cultures of *X. citri* were grown in 2xTY medium for 24 hours at 30 °C at 200 rpm and then maintained at room temperature (22 °C) without shaking for seven days. *X. citri* strains with knockouts in the genes for ΔX-Tfi^XA2610^, one of the X-T4SS components structural componentes (VirB4 to VirB10 and VirD4) and the double knockouts were used and are indicated above each well.

We then asked if specific individual components of the X-T4SS are necessary for the translocation of X-Tfe^XAC2609^ to the periplasm (Pathway (1) in Figure 1B). This is a a relevant hypothesis considering that cytoplasmic substrates of the T4SS are known to interact with various T4SS subcomplexes along the secretion pathway (Cascales and Christie 2004a; Atmakuri, Cascales, and Christie 2004; Cascales and Christie 2004b). To address this question, we deleted the genes coding for several X-T4SS subunits that are associated with the bacterial inner membrane (VirB4, VirB6, VirB8 and VirD4) (Macé et al. 2022), the outer membrane (VirB7, VirB9 and VirB10) (Fronzes, Christie, and Waksman 2009; Chandran et al. 2009; Souza et al. 2011; Oliveira et al. 2016; Sgro et al. 2018) and the VirB5 subunit believed to form part of the extracellular pilus (Alvarez-Martinez and Christie 2009; Christie, Whitaker, and González-Rivera 2014; Sheedlo et al. 2022). These deletions were introduced in both the *X. citri* wild-type and ΔX-Tfi^XAC2610^ genetic backgrounds. Growth of these strains in 24-well plates for 24 hours with agitation followed by five days without agitation revealed that all *X. citri* strains lacking X-Tfi^XAC2610^ cannot form biofilm, independent of the presence or absence of any X-T4SS structural component (Fig. 5B, **Fig. S5**). Taken together, these results show that, in the absence of X-Tfi^XAC2610^, *cis*-intoxication by X-Tfe^XAC2609^ inhibits biofilm formation in a manner that is independent of any subunit or subassembly of the X-T4SS apparatus.

Finally, as *X. citri* is the causal agent of citrus canker, we asked whether the absence of X-Tfi^XAC2610^ could affect the ability of this phytopathogen in causing disease. **Figure S6** shows that *X. citri* ΔX-Tfi^XAC2610^ can induce the appearance of citrus canker lesions in sweet orange leaves in a manner very similar to those caused by the wild-type strain. This phenotype was previously shown to be dependent on a type III secretion system (Cappelletti et al. 2011) but independent of the X-T4SS (Souza et al. 2011).

### Structural and co-evolutionary analyses of the X-Tfe^XAC2609^ - X-Tfi^XAC2610^ complex

Figure S7 shows the sequence conservation profile derived by the multiple sequence alignment of 429 non-redundant X-Tfi^XAC2610^ homologs (**Supplementary File 1**) and reveals several conserved motifs. One conserved motif is a region with several negatively charged amino acids corresponding to X-Tfi^XAC2610^ residues 151-159 that form a Ca^2+^-binding loop in the crystal structure of X-Tfi^XAC2610^ (Souza et al. 2015). **Figure S8** shows that the presence of Ca^2+^ ions significantly increases the thermal stability of X-Tfi^XAC2610^. This result suggests that the family of X-Tfi^XAC2610^ homologs could all be stabilized by divalent cation binding at this site.

To investigate the molecular mechanism of the immunity provided by X-Tfi^XAC2610^ against X-Tfe^XAC2609^ activity, we predicted the structure of the X-Tfe^XAC2609^/X-Tfi^XAC2610^ complex by AlphaFold2 (Mirdita et al. 2022; Varadi et al. 2022; Jumper et al. 2021) using the sequences of X-Tfi^XAC2610^ lacking the N-terminal signal peptide and the N-terminal GH10 glycohydrolase domain of X-Tfe^XAC2609^. Figure 6A shows that the best model of the X-Tfi^XAC2610^(54-267)/X-Tfe^XAC2609^(1-194) complex superposes with the previously determined crystal structure of X-Tfi^XAC2610^ (Souza et al. 2015). In this model, the interface between the two proteins involves another well-conserved motif in the X-Tfi^XAC2610^ family that includes a loop made up of residues 165-173 (Figure 6A-B and **Figure S9**). In the predicted X-Tfe^XAC2609^/X-Tfi^XAC2610^ complex, a highly conserved tyrosine (Y170) in this loop of X-Tfi^XAC2610^ directly interacts with the catalytic aspartate (E48) in the active site of X-Tfe^XAC2609^ (Fig. 6C, **Fig. S9** and **S10A**). To test the hypothesis that Y170 is involved in the mechanism of X-Tfi^XAC2610^-mediated inhibition of X-Tfe^XAC2609^, we carried out *in vitro* peptidoglycan hydrolysis assays using X-Tfe^XAC2609^ in the absence or presence of wild type X-Tfi^XAC2610^ or the X-Tfi^XAC2610^Y170A mutant. Figure 6D shows that the wild-type version of X-Tfi^XAC2610^ inhibits the hydrolytic activity of X-Tfe^XAC2609^, while the Y170A mutation strongly decreases peptidoglycan degradation.

**Figure 6.**
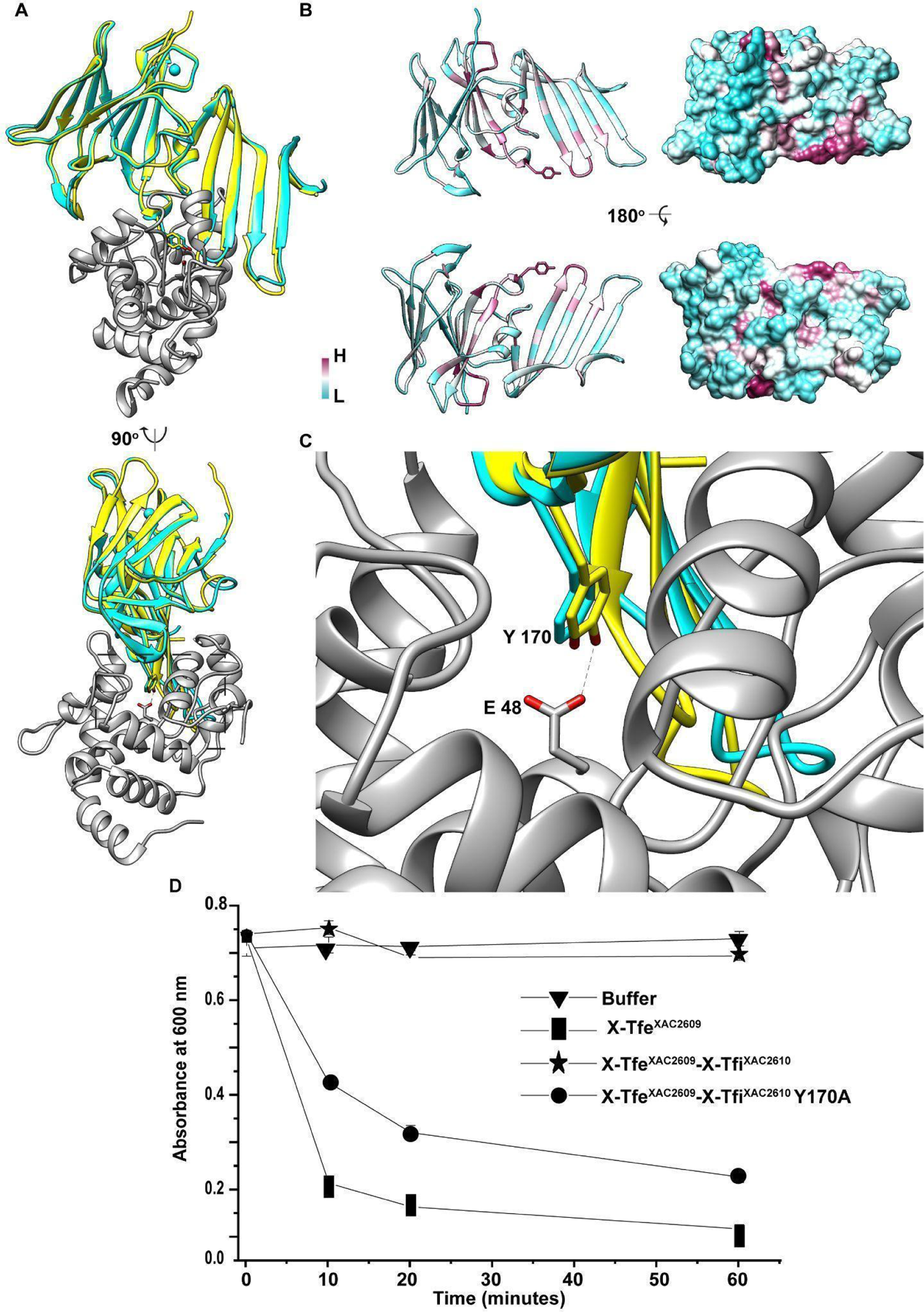
Co-evolutionary and structural analysis of the X-Tfe^XAC2609^-X-Tfi^XAC2610^ complex by AlphaFold2. **(A)** Ribbon representation of the X-Tfe^XAC2609^(1-194)-X-Tfi^XAC2610^(54-267) complex predicted by AlphaFold 2 (Mirdita et al. 2022; Varadi et al. 2022; Jumper et al. 2021). X-Tfe^XAC2609^(1-194) is shown in gray and X-Tfi^XAC2610^(54-267) is shown in yellow. The X-Tfi^XAC2610^(54-267) Alphafold2 model is superposed with the previously determined X-ray crystal structure of X-Tfi^XAC2610^ (residues 59-267, cyan; PDB code: 4QTQ) with a bound Ca^2+^ ion (sphere). Root-mean-square deviation (RMSD) of five Alphafold2 models is 0.22 Å, the Predicted Aligned Error (PAE) for the residues and the Predicted Local Distance Difference Test (lDDT) scores (**Figure S9**) also indicates that the models are internally consistent. **(B)** Cartoon (left) and surface (right) representations of X-Tfi^XAC2610^ colored according to degree of conservation (lowest, cyan; highest, purple). **(C)** Magnification of the protein-protein interface of the model shown in (A) that highlights the interaction between the conserved Y170 of X-Tfi^XAC2610^ and E48 of X-Tfe^XAC2609^ (both residues shown as sticks). **(D)** Peptidoglycan hydrolysis assay. *M. luteus* cell wall suspensions were treated with buffer (triangle) or with purified X-Tfe^XAC2609^(1-308) (square), X-Tfe^XAC2609^(1-308) - X-Tfi^XAC2610^(His-55–267) (star), X-Tfe^XAC2609^(1-308)-X-Tfi^XAC2610^(His-55–267) Y170A and absorbance was monitored at 650 nm. Error bars, s.d.; n= 3.

## Discussion

Peptidoglycan is a major component of the bacterial cell wall, functioning to maintain the stability of the cell envelope against turgor pressure (Callewaert et al. 2012; Silhavy, Kahne, and Walker 2010; Navarro et al. 2022). By far, the most well-studied PG hydrolase is lysozyme, which was the first natural antimicrobial molecule isolated from the human body (Fleming and Wright 1922). It targets the bacterial cell wall by hydrolyzing β(1→4) bonds between *N*-acetylmuramic acid and *N*-acetyl-D-glucosamine residues in the peptidoglycan layer (Lowe et al. 1967; Vocadlo et al. 2001). One century after its discovery, numerous studies have demonstrated a great diversity of lysozyme-like proteins that are widespread in all domains of life (Cernooka et al. 2022; Metcalf et al. 2014; Callewaert and Michiels 2010). Several families of lysozyme-like effectors are substrates of bacterial secretion systems in Gram-negative bacteria (Russell et al. 2012; Callewaert et al. 2012). Bacteria produce proteinaceous inhibitors in order to protect the PG from the activity of both endogenous and exogenous hydrolytic enzymes (Callewaert et al. 2012).

The lysozyme-like effector X-Tfe^XAC2609^ is a cytoplasmic protein that is transported in a X-T4SS-dependent manner into other bacterial cells. X-Tfi^XAC2610^, its cognate inhibitor, has a lipoprotein box motif for localization to the cell periplasm (Souza et al. 2015). We observed that cells expressing a functional X-Tfe^XAC2609^ in the absence of X-Tfi^XAC2610^ completely abolished bacterial biofilm formation and that biofilms are still not formed by cells with this genetic background when essential X-T4SS components are knocked out. These results show for the first time that inhibition of PG hydrolases by immunity proteins can be required for bacterial biofilm formation. Accordingly, another example of the relationship of PG stability and biofilm formation was described in *Campylobacter jejuni*, where PG acetylation is associated with the maintenance of cell wall integrity and contributes to biofilm establishment (Iwata et al. 2016). Also, inhibition of peptidoglycan transpeptidation and transglycosylation by D-Leucine and flavomycin can specifically impair biofilm formation in Gram-positive bacteria, although with minimal impact on the bacterial planktonic lifestyle (Bucher et al. 2015). These studies corroborate our results that X-Tfe^XAC2609^-induced damage to the cell envelope compromises biofilm formation.

Two general, not necessarily exclusive, hypotheses were proposed to account for X-Tfe^XAC2609^-dependent toxicity in the absence of X-Tfi^XAC2610^. The first, which we call *trans*-intoxication or fratricide, hypothesizes that *X. citri* cells inject a cocktail of X-Tfes, including X-Tfe^XAC2609^, into neighboring *X. citri* cells and, in the absence of at least one cognate immunity protein, could cause cellular injury and consequent growth suppression or cell death. *Trans*-intoxication has been shown to occur in *P. aeruginosa* cells via the H1-T6SS-dependent secretion of the PG hydrolases Tse1 and Tse3 (Russell et al., 2011). Also, in *Vibrio cholerae*, T6SS-associated immunity proteins have been shown to be important in preventing *trans*-intoxication (Dong et al. 2013). However, in the case of the X-Tfe^XAC2609^-dependent toxicity observed in the absence of X-Tfi^XAC2610^, a *trans*-intoxication mechanism is not consistent with the observation that both the detrimental effect of X-Tfe^XAC2609^ and protection conferred by X-Tfi^XAC2610^ are independent of a functional X-T4SS and that wild-type *X. citri* does not outgrow *X. citri* strains lacking X-Tfi^XAC2610^ or several other immunity proteins in competition assays (Figure 2B **and movie S7**). Furthermore, X-Tfe^XAC2609^-dependent autolysis in the absence of X-Tfi^XAC2610^ does not require the former’s C-terminal XVIPCD X-T4SS secretion signal. Thus, we can discard the *trans*-intoxication mechanism or at least propose that this pathway contributes only a small fraction of the X-Tfe^XAC2609^-dependent toxic effects observed in the absence of X-Tfi^XAC2610^. Nevertheless, the fact that wild-type *X. citri* is unable to kill strains lacking immunity proteins is intriguing. That cells in some way avoid *trans*-intoxication is revealed by the fact that *X. citri* wild-type cells carrying an X-T4SS and full cohort of X-Tfes do not kill the *X. citri Δ8Δ2609-GFP*, the *X. citri* ΔX-Tfe^XAC2609^ΔX-Tfi^XAC2610^, or any other X-T4SS-deficient strain tested points to a still-to-be-characterized mechanism of protection against *trans*-intoxication (fratricide) that will be addressed in future studies by our group.

We are thus left to consider the *cis*-intoxication hypothesis to explain the toxicity of X-Tfe^XAC2609^ and the protection conferred by X-Tfi^XAC2610^. Here, the effector exerts its toxic effects within the cell in which it was synthesized. Two cytoplasmic *P. aeruginosa* T6SS effectors, Tse2 and Tse6, have been shown to cause cis-intoxication in a T6SS null strain when immunity is depleted (M. Li et al. 2012; Whitney et al. 2015). In the case of X-Tfe^XAC2609^, the toxin somehow makes its way into the cell periplasm where, in the absence of X-Tfi^XAC2610^, it degrades the peptidoglycan layer. Analysis of the X-Tfe^XAC2609^ sequence by the SignalP 6.0 (Teufel et al. 2022) and other algorithms failed to detect any putative N-terminal signal peptide. Although the mechanism responsible for X-Tfe^XAC2609^ transfer into the periplasm is at the moment unknown, we have shown that it is independent of a functional X-T4SS and of the XVIPCD secretion signal. Other bacterial proteins have been shown to transfer into the periplasm without any obvious secretion signal, for example VgrG3 from *Vibrio cholerae* (Ho et al. 2017), CI2 and in HdeA expresssed in *E.coli* (Barnes and Pielak 2011).

We have previously pointed out that the structure of X-Tfi^XAC2610^ adopts a similar β-propeller fold topology to two other peptidoglycan hydrolase inhibitors: the *P. aeruginosa* Type VI immunity protein Tsi1 and the *Aeromonas hydrophila* periplasmic i-type lysozyme inhibitor PliI-Ah (Souza et al. 2015). Interestingly, both of these inhibitors block the activity of their respective targets by inserting an exposed loop into the enzyme’s active site, in a manner similar to that predicted for the X-Tfe^XAC2609^-X-Tfi^XAC2610^ complex (**Fig. S10**); this in spite of the fact that they share very low sequence identity with X-Tfi^XAC2610^ (7% for Tsi1 and 12% for PliI-Ah; (Souza et al. 2015). In the case of PliI-Ah, its crystal structure in complex with the i-type lysozyme from *Meretrix lusoria* (Ml-iLys) revealed a complementary key-lock interface through the interaction of an exposed loop of PliI-Ah into the substrate-binding groove of Ml-iLys (Herreweghe et al. 2015). Comparison of the topology diagrams of X-Tfi^XAC2610^, PliI-Ah and Tsi1 shows that the loops they use to interact with their cognate targets are topologically equivalent (**Fig. S11**). While X-Tfi^XAC2610^ is predicted to use Y170 to interact directly with E48 in the active site of X-Tfe^XAC2609^, in Tsi1 a serine residue (S109) of its inhibitory insertion loop directly interacts with H91 in the active site of the target Tse1 PG amidase enzyme (Benz et al. 2012) (**Fig. S10**). In a similar manner, in PliI-Ah, a serine residue (S104) directly recognizes the active site residue E18 of the inhibited lysozyme (Herreweghe et al. 2015) (**Fig. S10**).

The structural similarities in the inhibitory mechanisms of X-Tfi^XAC2610^, Tsi1 and PliI-Ah are intriguing. While X-Tfi^XAC2610^ and PliI-Ah inhibit PG glycohydrolases, Tsi1 inhibits the PG amidase activity of Tse1. It is also worthy to note that whereas X-Tfi^XAC2610^ inhibits the *cis*-intoxication activity of its cognate effector, the biological function of Tsi1 was described to protect *P. aeruginosa* cells from the deleterious effects of Tse1 molecules transferred from neighboring cells via the H1-T6SS, thus avoiding fratricide or *trans*-intoxication (Russell et al. 2011). Therefore, there are interesting structural and functional relationships among PG hydrolase inhibitors from distinct and diverse biological systems and, even though X-Tfi^XAC2610^, Tsi1 and PliI-Ah do not share any detectably relevant sequence identity, they may in fact be very distant members of a homologous protein superfamily that have evolved to inhibit PG hydrolases (such as glycohydrolases and amidases).

## Acknowledgments

We thank Dr. Frederico Gueiros-Filho for support in the microscopy studies, Central Analítica at Instituto de Química, Universidade de São Paulo and Dr. Ii-Sei Watanabe and Dr. Sonia R. Yokomizo from the Instituto de Ciências Biológicas, Universidade de São Paulo for their support and assistance in transmission electron microscopy sample preparation and image acquisition. This work was supported by grants from the Fundação de Amparo à Pesquisa do Estado de São Paulo (FAPESP) to C.S.F. (Projects #2017/17303-7 and #2021/10577-0) and FAPESP fellowships to G.U.O (Project #2018/09277-9) and D.P.S. (Project #2011/50521-1). D.P.S. and G.U.O. acknowledge CAPES and CNPq fellowships.

## SUPPORTING INFORMATION

**S1 Table.**
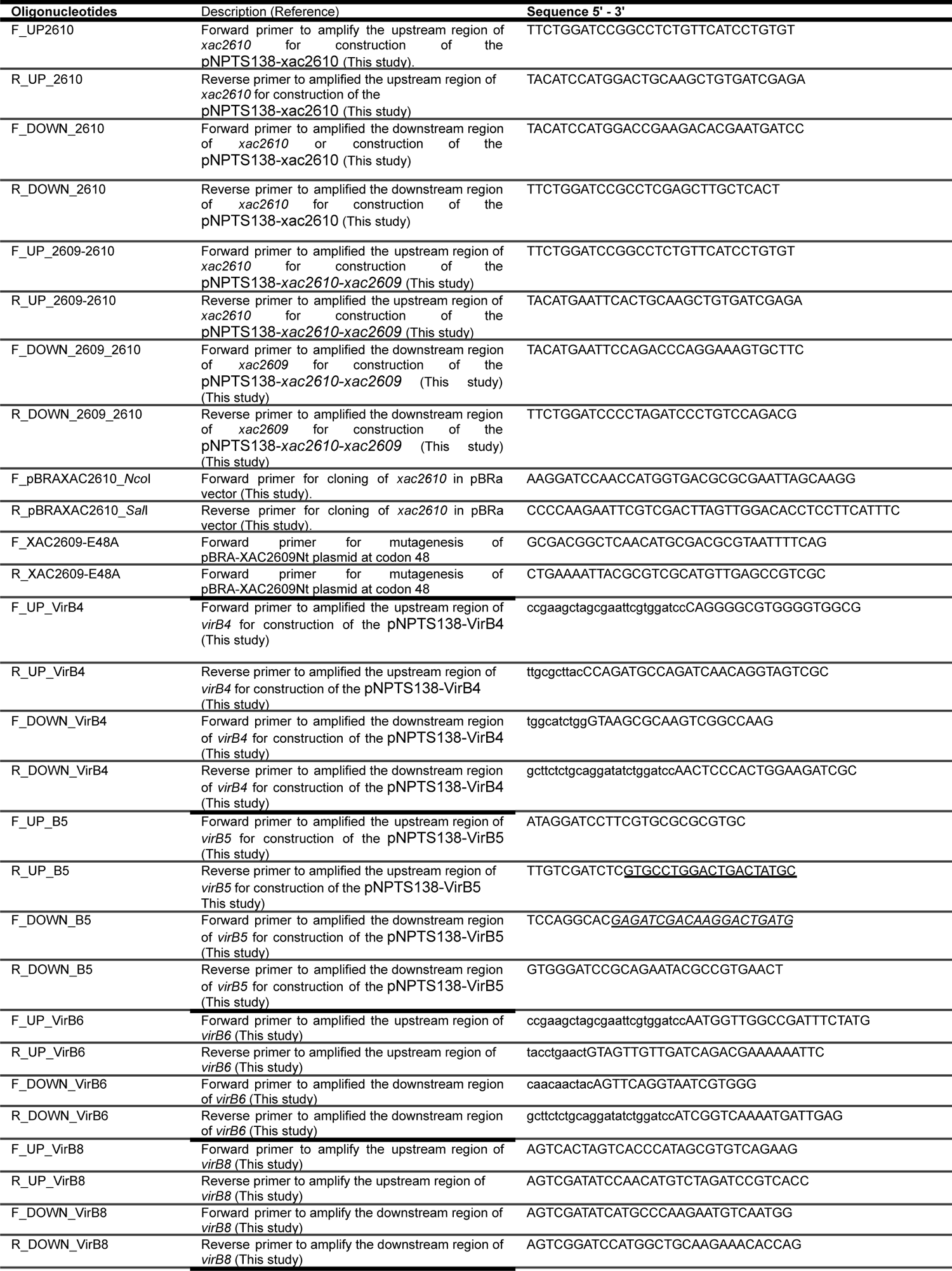
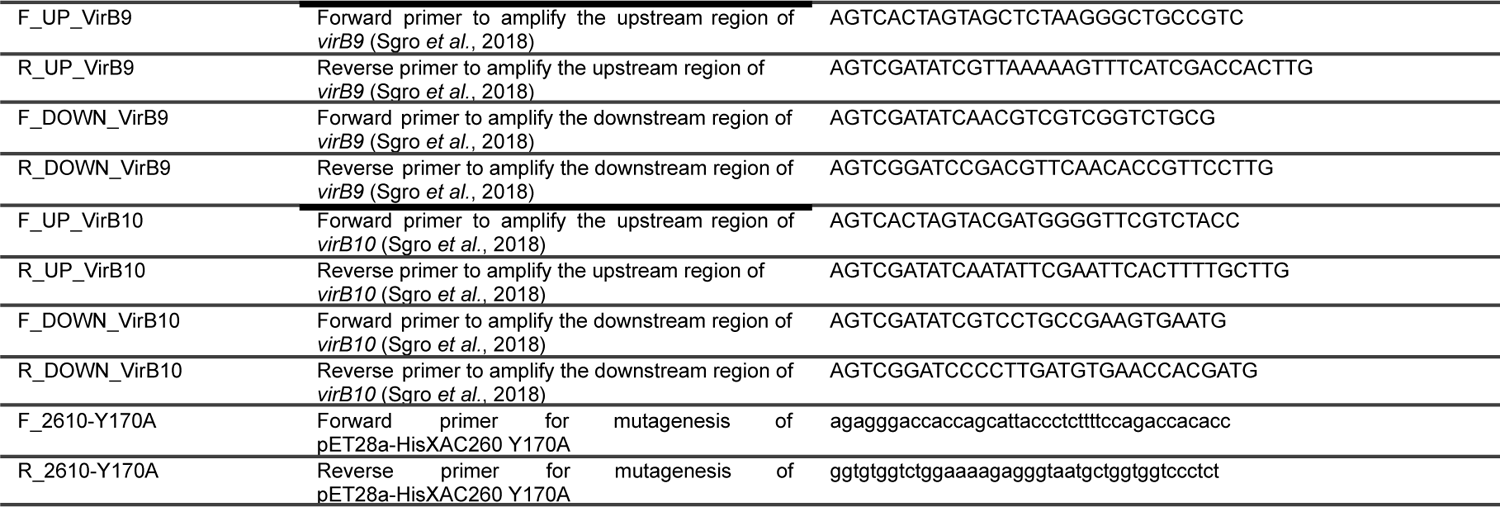
Oligonucleotides used in this study.

**S2 Table.** Plasmids used in this study.

| Plasmids | Properties | References |
| --- | --- | --- |
| **pBRA** | pBAD24 derivative vector which has an arabinose-inducible promoter that is constitutively active plasmid in *Xac*; Sp<sup>r</sup> | M.Marroquin,unpublished |
| **pBRA-XAC2609Nt** | pBRA encoding X-Tfe<sup>XAC2609</sup> N-terminus (residues 1-306); Sp<sup>r</sup> | (Souza et al. 2015) |
| **pBRA-XAC2609NtE48A** | pBRA-XAC2609Nt with a point mutation at codon 48; Sp<sup>r</sup> | This study |
| **pBRA-2609FL** | pBRA encoding X-Tfe<sup>XAC2609</sup> full length (residues 1-431); Sp<sup>r</sup> | (Souza et al. 2015) |
| **pBRA-XAC2610** | pBRA encoding X-Tfi<sup>XAC2610</sup>; Sp<sup>r</sup> | This study |
| **pBRA-VirB7** | pBRA encoding VirB7 | (Souza et al. 2015) |
| **pNPTS138** | Suicide vector for generation of gene knockouts (*sacB*), Km<sup>r</sup> | M.R.Alley unpublished |
| **pNPTS138-xac2610** | Suicide vector for generation of the *X. citri* ΔX-Tfi<sup>XAC2610</sup>, and doubles deletion of ΔX-Tfi<sup>XAC2610</sup> and ΔVir(B4-B10,D4) ΔX-Tfi<sup>XAC2610</sup> strains. | This study |
| **pNPTS138-xac2610-xac2609** | Suicide vector for generation of the *X. citri*ΔX-Tfe<sup>XAC2609</sup> ΔX-Tfi<sup>XAC2610</sup> strain; Km<sup>r</sup> | This study |
| **pBBR-5GFP** | Broad host range plasmid cloning vector pBBR1MCS-5 expressing GFP; Gm<sup>r</sup> | (Dunger et al. 2014) |
| **pBBR-2RFP** | Broad host range plasmid cloning vector pBBR1MCS-5 expressing RFP; Gm<sup>r</sup> | (Oka et al. 2022) |
| **pET28a-XAC2610His-22-267** | Production of the His-tagged HIS-X-Tfi<sup>XAC2610</sup>(22-267) | (Alegria et al. 2005) |
| **pET28a-XAC2610His-22-267Y170A** | Production of the His-tagged HIS-X-Tfi<sup>XAC2610</sup>(22-267) Y170A | This study |
| **pET11a-XAC2609(1-308)** | Eptopic Production of the X-Tfe<sup>XAC2609</sup>(1-308) | (Souza et al. 2015) |
| **pET11a-XAC2610(55-267)** | Production of the XTfi<sup>XAC2610</sup> folded domain (residues 55-267) | (Souza et al. 2015) |
| **pNPTS138-VirB4** | Suicide vector for deletion of *virB4* in the *X. citri* ΔVirB4, Km<sup>r</sup> | This study |
| **pNPTS138-VirB5** | Suicide vector for deletion of *virB5* in the *X. citri* ΔVirB5, Km<sup>r</sup> | This study |
| **pNPTS138-VirB6** | Suicide vector for deletion of *virB6* in the *X. citri* ΔVirB6, Km<sup>r</sup> | This study |
| **pNPTS138-VirB7** | Suicide vector for deletion of *virB7* in the *X. citri* in ΔVirB7 Δ/ Km<sup>r</sup> | (Souza et al. 2011) |
| **pNPTS138-VirB8** | Suicide vector for deletion of *virB8* in *X. citri* ΔVirB8, Km<sup>r</sup> | This study |
| **pNPTS138-VirB9** | Suicide vector for deletion of *virB9* in *X. citri* ΔVirB9, Km<sup>r</sup> | (Sgro et al. 2018) |
| **pNPTS138-VirB10** | Suicide vector for deletion of *virB10* in *X. citri* ΔVirB10 Km<sup>r</sup> | (Sgro et al. 2018) |
| **pNPTS138-VirD4** | Suicide vector for deletion of *virD4* in *X. citri* ΔVirD4, Km<sup>r</sup> | (Souza et al, 2015) |

**S3 Table.** Bacterial strains used in this study.

| Bacterial strains | Genotype | reference |
| --- | --- | --- |
| *X. citri* wild type | *Xanthomonas citri* subsp. *citri* strain 306, wild type strain; Ap<sup>r</sup> | (da Silva et al. 2002) |
| *X. citri* Δ*virB7* | *X. citri* Δ*virB7* XAC2622; Ap<sup>r</sup> | (Souza et al. 2015) |
| *X. citri* ΔX-Tfi<sup>XAC2610</sup> | *X. citri* Δ*xac2610*, Ap<sup>r</sup> | This study |
| *X. citri* wild type pBRA GFP | *X. citri* wild type pBRA pBBR-5GFP; Ap<sup>r</sup> Gm<sup>r</sup> | (Souza et al. 2015) |
| *X. citri* Δ*virD4* GFP | *X. citri* Δ*virD4*XAC2623 pBBR-5GFP; Ap<sup>r</sup> | (Souza et al. 2015) |
| *X. citri* ΔX-Tfi<sup>XAC2610</sup> pBRA | *X. citri* Δ*xac2610* pBRA; Ap<sup>r</sup> Sp<sup>r</sup> | This study |
| *X. citri* ΔX-Tfi<sup>XAC2610</sup> pBRA GFP | *X. citri* Δ*xac2610* pBRA pBBR-5GFP; Ap<sup>r</sup> Sp<sup>r</sup> Gm<sup>r</sup> | This study |
| *X. citri* ΔX-Tfi<sup>XAC2610</sup> + X-Tfi<sup>XAC2610</sup> | *X. citri* Δ*xac2610* pBRA-XAC2610; Ap<sup>r</sup> Sp<sup>r</sup> | This study |
| *X. citri* ΔX-Tfi<sup>XAC2610</sup> + X-Tfi<sup>XAC2610</sup> GFP | *X. citri* Δ*xac2610* pBRA-XAC2610 pBBR-5GFP; Ap<sup>r</sup> Sp<sup>r</sup> Gm<sup>r</sup> | This study |
| *X. citri* ΔX-Tfi<sup>XAC2610</sup> Δ*virB7* | *X. citri* Δ*virB7*XAC2622 Δ*xac2610*; Ap<sup>r</sup> | This study |
| *X. citri* ΔX-Tfi<sup>XAC2610</sup> Δ*virB7* pBRA | *X. citri* Δ*virB7*XAC2622 Δ*xac2610* pBRA; Ap<sup>r</sup> Sp<sup>r</sup> | This study |
| *X. citri* ΔX-Tfi<sup>XAC2610</sup> Δ*virB7* pBRA GFP | *X. citri* Δ*virB7*XAC2622 Δ*xac2610* pBRA pBBR-5GFP; Ap<sup>r</sup> Sp<sup>r</sup> Gm<sup>r</sup> | This study |
| *X. citri* ΔX-Tfi<sup>XAC2610</sup> Δ*virB7* + X-Tfi<sup>XAC2610</sup> | *X. citri* Δ*virB7*XAC2622 Δ*xac2610* pBRA-XAC2610; Ap<sup>r</sup> Sp<sup>r</sup> | This study |
| *X. citri* ΔX-Tfi<sup>XAC2610</sup> Δ*virB7* + X-Tfi<sup>XAC2610</sup> GFP | *X. citri* Δ*virB7*XAC2622 Δ*xac2610* pBRA-XAC2610 pBBR-5GFP; Ap<sup>r</sup> Sp<sup>r</sup> Gm<sup>r</sup> | This study |
| *X. citri* ΔX-Tfi<sup>XAC2610</sup> Δ*virD4* GFP | *X. citri* Δ*virD4*XAC2623 Δ*xac2610* pBBR-5GFP; Ap<sup>r</sup> Gm<sup>r</sup> | This study |
| *X. citri* ΔX-Tfi<sup>XAC2610</sup> ΔTfe<sup>XAC2609</sup> | *X. citri* Δ*xac2610*Δ*xac2609*; Ap<sup>r</sup> | This study |
| *X. citri* ΔX-Tfe<sup>XAC2609</sup> ΔX-Tfi<sup>XAC2610</sup> pBRA | *X. citri* Δ*xac2610*Δ*xac2609* pBRA; Ap<sup>r</sup> Sp<sup>r</sup> | This study |
| *X. citri* ΔX-Tfi<sup>XAC2610</sup> ΔX-Tfe<sup>XAC2609</sup> pBRA GFP | *X. citri* Δ*xac2610*Δ*xac2609* pBRA pBBR-5GFP; Ap<sup>r</sup> Sp<sup>r</sup> Gm<sup>r</sup> | This study |
| *X. citri* ΔX-Tfe<sup>XAC2609</sup> ΔX-Tfi<sup>XAC2610</sup> + X-Tfe<sup>XAC2609</sup>FL GFP | *X. citri* Δ*xac2610*Δ*xac2609* pBRA-2609FL pBBR-5GFP; Ap<sup>r</sup> Sp<sup>r</sup> Gm<sup>r</sup> | This study |
| *X. citri* ΔX-Tfe<sup>XAC2609</sup> ΔX-Tfi<sup>XAC2610</sup> + X-Tfe<sup>XAC2609</sup>NT GFP | *X. citri* Δ*xac2610*Δ*xac2609* pBRA-2609NT pBBR-5GFP; Ap<sup>r</sup> Sp<sup>r</sup> Gm<sup>r</sup> | This study |
| *X. citri* ΔX-Tfe<sup>XAC2609</sup> ΔX-Tfi<sup>XAC2610</sup> + X-Tfe<sup>XAC2609</sup>NT E48A GFP | *X. citri* Δ*xac2610*Δ*xac2609* pBRA-XAC2609NtE48A pBBR-5GFP; Ap<sup>r</sup> Sp<sup>r</sup> Gm<sup>r</sup> | This study |
| *X. citri* Δ*virB4* | *X. citri* Δ*virB4* XAC2614; Ap<sup>r</sup> | This study |
| *X. citri* Δ*virB5* | *X. citri* Δ*virB5* XAC2613; Ap<sup>r</sup> | This study |
| *X. citri* Δ*virB6* | *X. citri* Δ*virB6* XAC2612; Ap<sup>r</sup> | This study |
| *X. citri* Δ*virB8* | *X. citri* Δ*virB8* XAC2621; Ap<sup>r</sup> | (Sgro et al. 2018) |
| *X. citri* Δ*virB9* | *X. citri* Δ*virB9* XAC2620; Ap<sup>r</sup> | (Sgro et al. 2018) |
| *X. citri* Δ*virB10* | *X. citri* Δ*virB10* XAC2619; Ap<sup>r</sup> | (Sgro et al. 2018) |
| *X. citri* ΔX-Tfi<sup>XAC2610</sup> Δ*virB4* | *X. citri* Δ*virB4* XAC2614; Δ*xac2610*; Ap<sup>r</sup> | This study |
| *X. citri* ΔX-Tfi<sup>XAC2610</sup> Δ*virB5* | *X. citri* Δ*virB5* XAC2613; Δ*xac2610*; Ap<sup>r</sup> | This study |
| *X. citri* ΔX-Tfi<sup>XAC2610</sup> Δ*virB6* | *X. citri* Δ*virB6* XAC2612; Δ*xac2610* Ap<sup>r</sup> | This study |
| *X. citri* ΔX-Tfi<sup>XAC2610</sup> Δ*virB8* | *X. citri* Δ*virB8* XAC2621; Δ*xac2610*; Ap<sup>r</sup> | This study |
| *X. citri* ΔX-Tfi<sup>XAC2610</sup> Δ*virB9* | *X. citri* Δ*virB9* XAC2620; Δ*xac2610*; Ap<sup>r</sup> | This study |
| *X. citri* ΔX-Tfi<sup>XAC2610</sup> Δ*virB10* | *X. citri* Δ*virB10* XAC2619; Δ*xac2610*; Ap<sup>r</sup> | This study |
| *E. coli* MG1655 |  |  |
| *X. citri* Δ8Δ2609-GFP | *X. citri* strain strain deleted of the *X-Tfe*/i<sup>XAC2885/XAC2884, *X-Tfe*/i<sup>XAC0574/XAC0573, *X-Tfe*/i<sup>XAC0096/XAC0097, *X-Tfe*/i<sup>XAC3634/XAC3633, *X-Tfe*/i<sup>XAC0466/XAC047, *X-Tfe*/i<sup>XAC3266/XAC3267, *X-Tfe*/i<sup>XAC4264/XAC42633/XAC4262 with *X-Tfe*<sup>XAC2609</sup> substituted by msGFP; Ap<sup>r</sup></sup></sup></sup></sup></sup></sup></sup> | (Oka et al. 2022) |
| *X. citri* Δ8Δ2609-GFP kan<sup>R</sup> | *X. citri* strain Δ8Δ2609-GFP+ pBBR-2RFP; Ap<sup>R</sup>, Km<sup>R</sup> | This study |
| *X. citri* ΔX-Tfe<sup>XAC2609</sup> ΔX-Tfi<sup>XAC2610</sup> kan<sup>R</sup> | *X. citri* ΔX-Tfe<sup>XAC2609</sup> ΔX-Tfi<sup>XAC2610</sup> + pBBR-2RFP; Ap<sup>R</sup>, kan<sup>R</sup> | This study |
| *X. citri* Δ*virB7* kan<sup>R</sup> | *X. citri* Δ*virB7* + pBBR-2RFP; Ap<sup>R</sup>, kan<sup>R</sup> | This study |
| *X. citri* wild-type kan<sup>R</sup> | *X. citri* wild-type + pBBR-2RFP; Ap<sup>R</sup>, kan<sup>R</sup> | This study |
| *X. citri* wild-type gent<sup>R</sup> | *X. citri* Wild-type + pBBR-5GFP; Ap<sup>R</sup>, kan<sup>R</sup> | This study |

**S4 Table.**
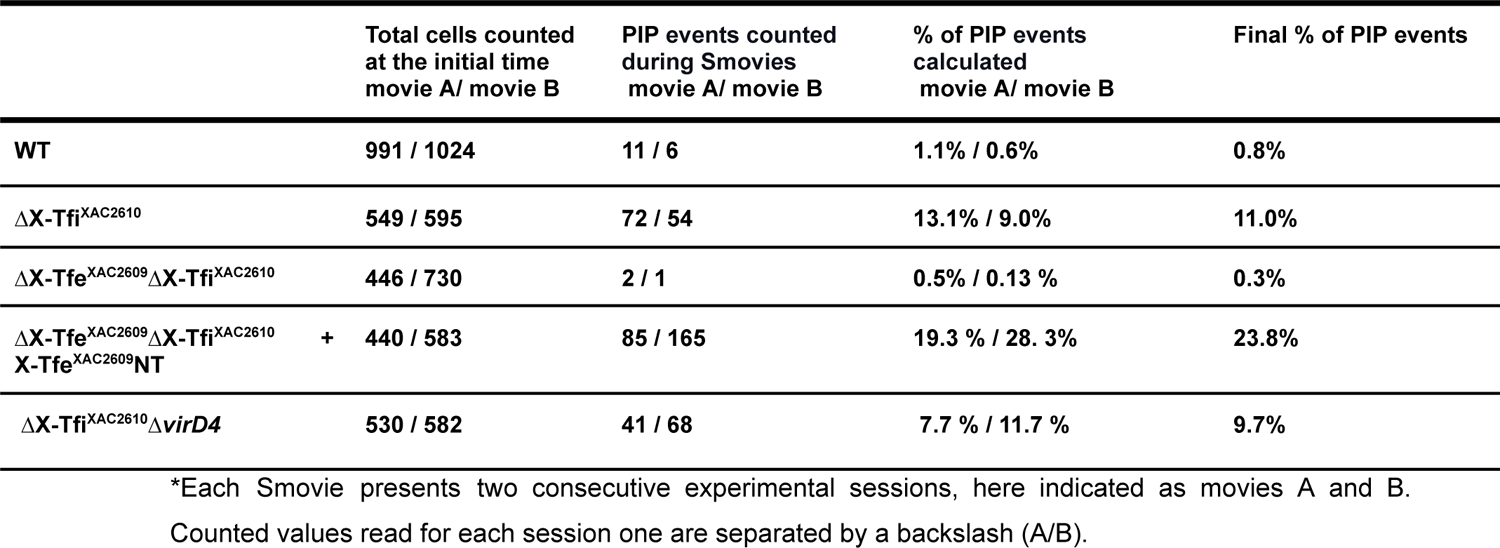
Analysis of cellular propidium iodide permeability (PIP) in Smovies 1-5*.

**S5 Table.** Analysis TEM micrographs of *X. citri* strains.

| | WT | $\Delta virB7$ | $\Delta X-Tfi^{XAC2610}$ | $\Delta X-Tfi^{XAC2610} C$ | $\Delta X-Tfi^{XAC2610}$<br>$\Delta VirB7$ |
| --- | --- | --- | --- | --- | --- |
| total number of cells<br>counted | 2640 | 675 | 2152 | 1212 | 1337 |
| micrographs<br>analyzed | 15 | 5 | 10 | 8 | 10 |
| *mean of cells per<br>micrograph | 176 +/- 11.7 | 135 +/- 20.9 | 215 +/- 17.9 | 151 +/- 9.8 | 166 +/- 7.8 |
| total of damaged<br>cells identified | 3 | 1 | 39 | 4 | 15 |
| mean of damaged<br>cells per micrograph | 0.2 +/- 0.1 | 0.2 +/- 0.2 | 3.9 +/- 0.76 | 0.5 +/- 0.5 | 1.5 +/- 0.45 |
| percentage of<br>damaged cells per<br>micrograph | 0.11% +/- 0.06% | 0.15% +/- 0.15% | 1.95% +/- 0.39% | 0.4% +/- 0.4% | 0.88% +/- 0.25% |

**Figure S1.**
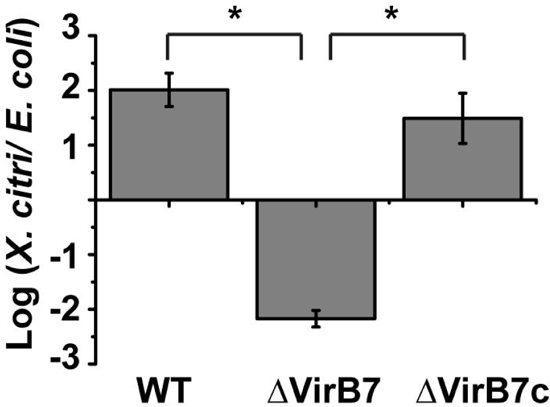
Inter-species bacterial competition assay. Ratio of the number of viable *X. citri* and *E. coli* Bl21(DE3) cells after 48 hours of co-culture on Luria-Bertani (LB) agar at 28 °C. Asterisks (*) indicate significant differences at the *p-value* <0.001 (ANOVA). Error bars ± s.d. n=4. *X. citri* strains: WT (wild type carrying the empty pBRA vector), ΔVirB7 (*virB7* knockout strain carrying the empty pBRA vector), ΔVirB7c (*virB7* knockout carrying the plasmid pBRA-VirB7).

**Figure S2.**
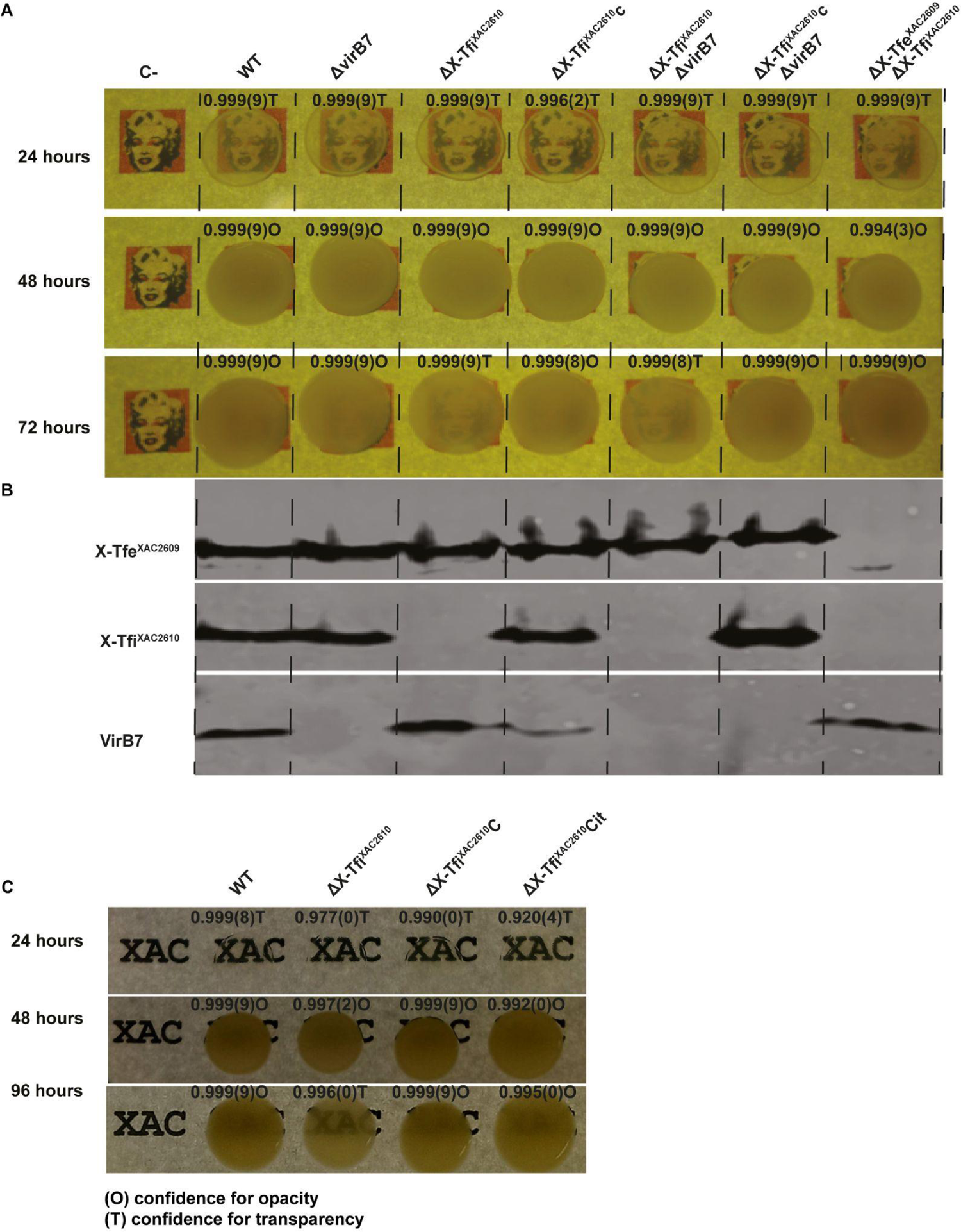
Colony opacity assay. **(A)** *X. citri* wild-type mutant strains were grown on an LB agar plate placed above a paper sheet printed with a picture. Plate images were acquired after 24, 48 and 72 hours of growth at 28°C. *X. citri* wild-type (WT) strain, ΔVirB7 strain, ΔX-Tfi^XAC2610^ strain, ΔX-Tfi^XAC2610^ΔVirB7 strain, ΔX-Tfe^XAC2609^ΔX-Tfi^XAC2610^ strain, and complemented strains (ΔX-Tfi^XAC2610^ + X-Tfi^XAC2610^ (ΔX-Tfi^XAC2610^c), ΔX-Tfi^XAC2610^ΔVirB7 + X-Tfi^XAC2610^ (ΔX-Tfi^XAC2610^cΔVirB7) used are indicated above each lane. Above each colony are the convolutional neural network confidence tendency index for opacity (O) and transparency (T) as described in Materials and Methods. **(B)** Immunodetection by Western Blot of X-Tfe^XAC2609^, X-Tfi^XAC2610^, and VirB7 of the cell extract of each *X. citri* strain used in (A). **(C)** *X. citri* wild-type and derived mutants were grown on an LB agar plate placed above a paper sheet printed with the letters “XAC”. Plate images were acquired after 24, 48, and 96 hours of growth. *X. citri* wild-type (WT), ΔX-Tfi^XAC2610^, ΔX-Tfi^XAC2610^ transformed with pBRA-X-Tfi^XAC2610^ (ΔX-Tfi^XAC2610^c), ΔX-Tfi^XAC2610^ transformed with pBRA-X-Tfi^XAC2610^His-22-267 (ΔX-Tfi^XAC2610^cit).

**Figure S3.**
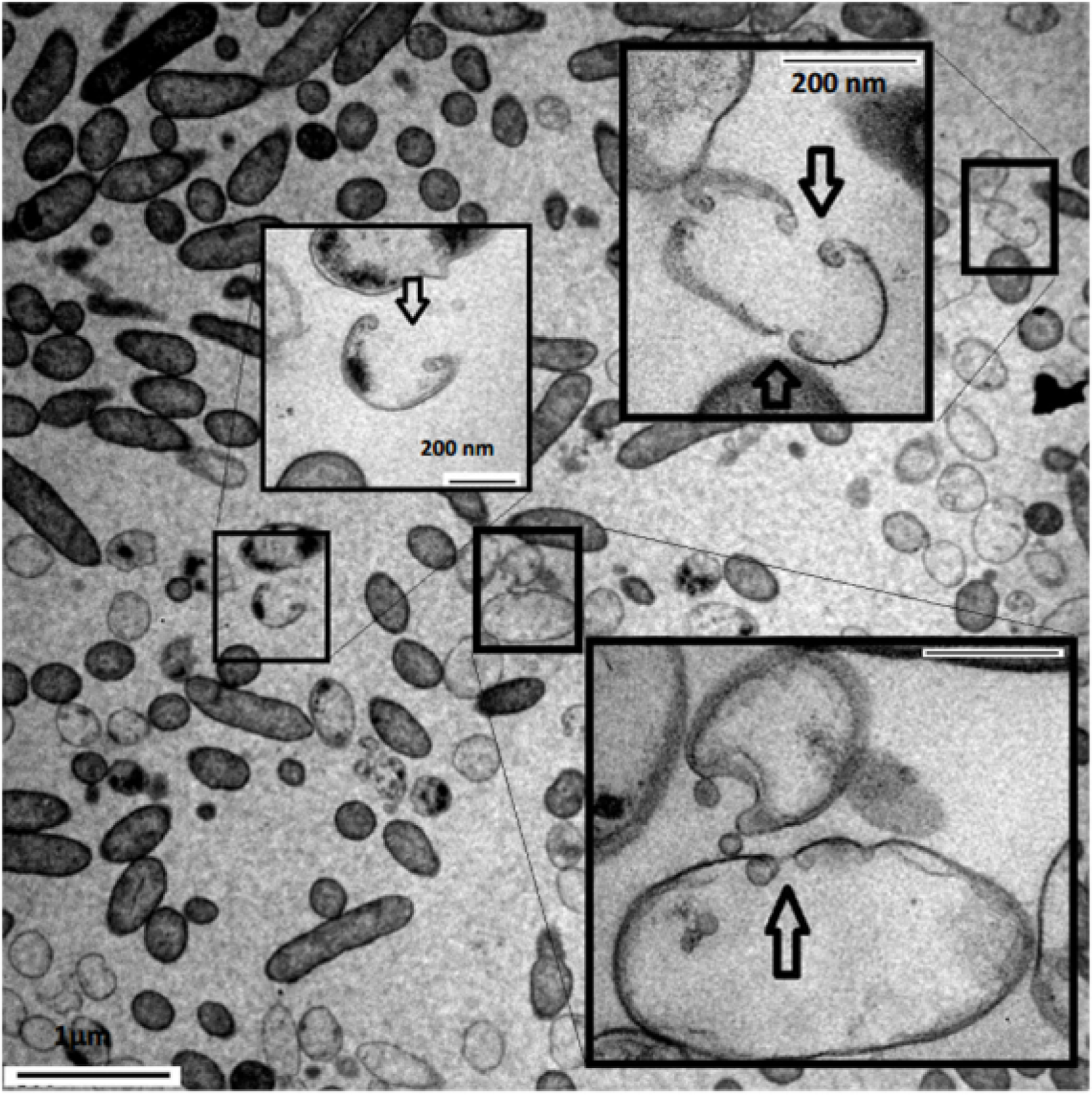
Transmission electron microscopy (TEM) of *X. citri* ΔX-Tfi^XAC2610^ cells. Insets highlight cells having an impaired enveloped cellular (arrows).

**Figure S4.**
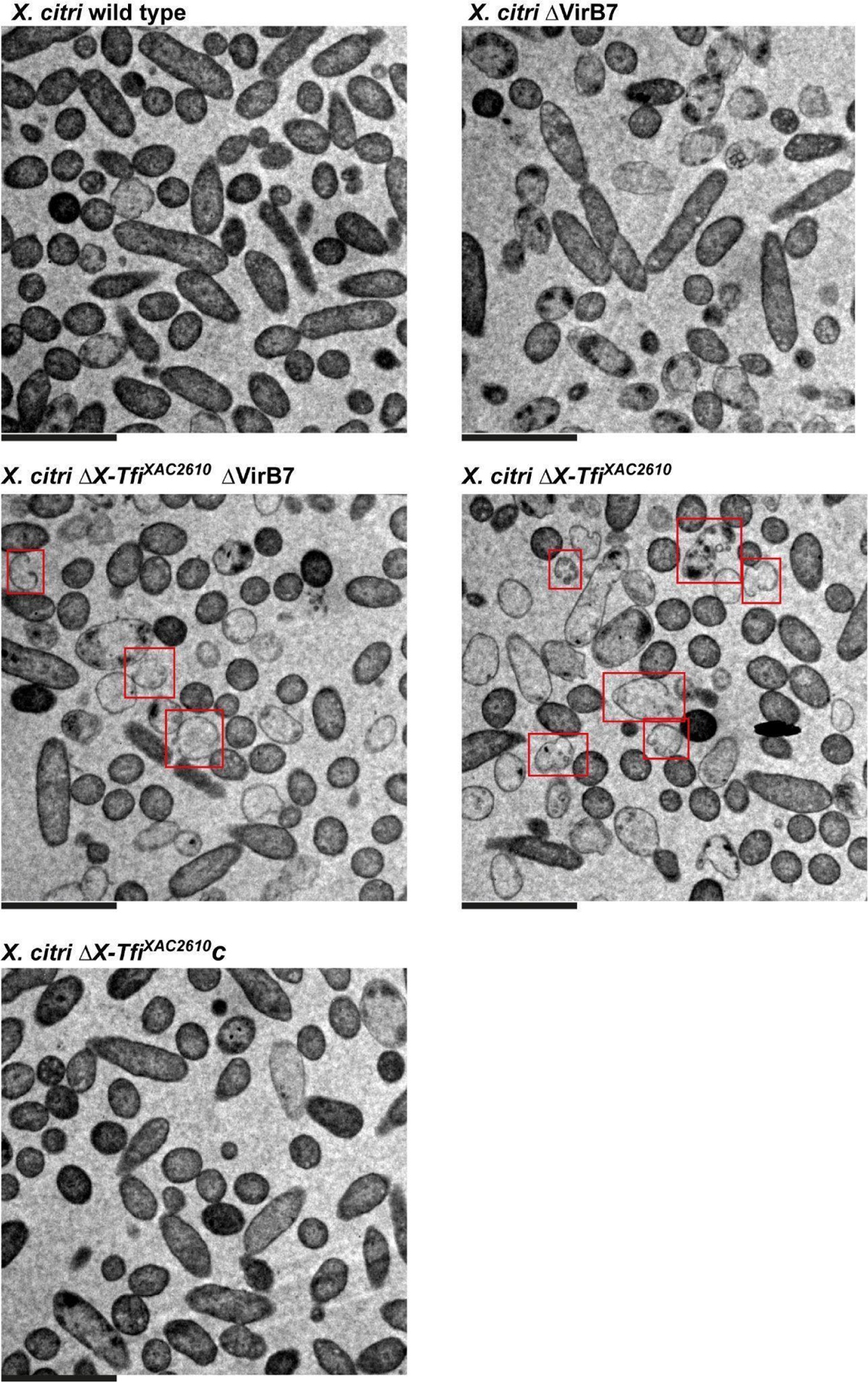
Transmission electron microscopy (TEM) of different X. citri strains. Micrographs of *X. citri* wild-type, ΔX-Tfi^XAC2610^, ΔVirB7, ΔX-Tfi^XAC2610^-ΔVirB7 and ΔX-Tfi^XAC2610^ complemented with pBRA-XAC2610 vector (ΔX-Tfi^XAC2610^c). Examples of damaged cells are highlighted in red rectangles. Scale bar = 1 µm.

**Figure S5.**
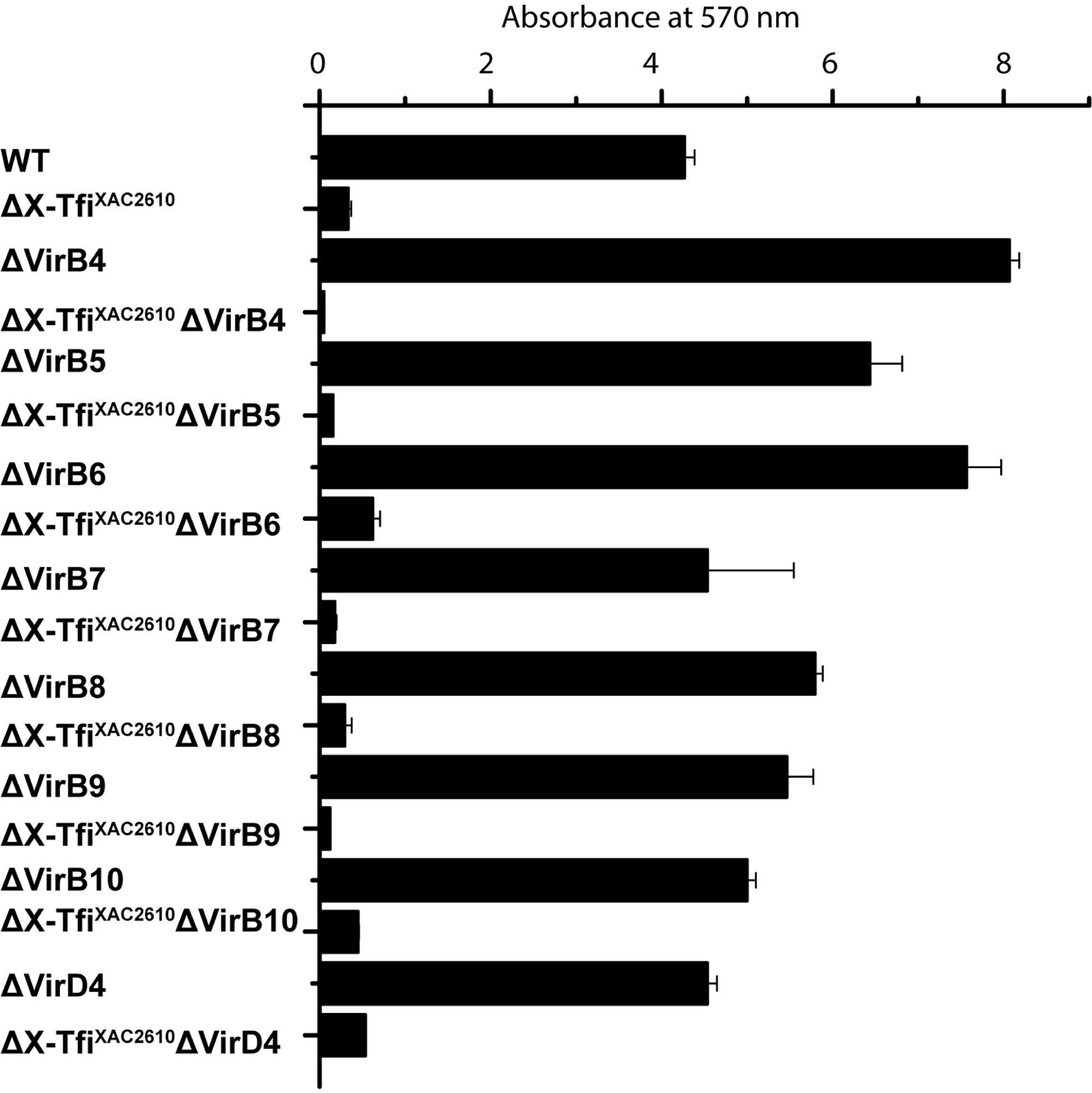
*X. citri* biofilm quantification using crystal violet. *X. citri* strains used in the assay are indicated in the figure legend. *X. citri* cells were grown using 24 well-plate (e.g., Fig. 4B) in 2xTY media for seven days of cultivation at 22 °C. Error bars, s.d. n= 3 experiments.

**Figure S6.**
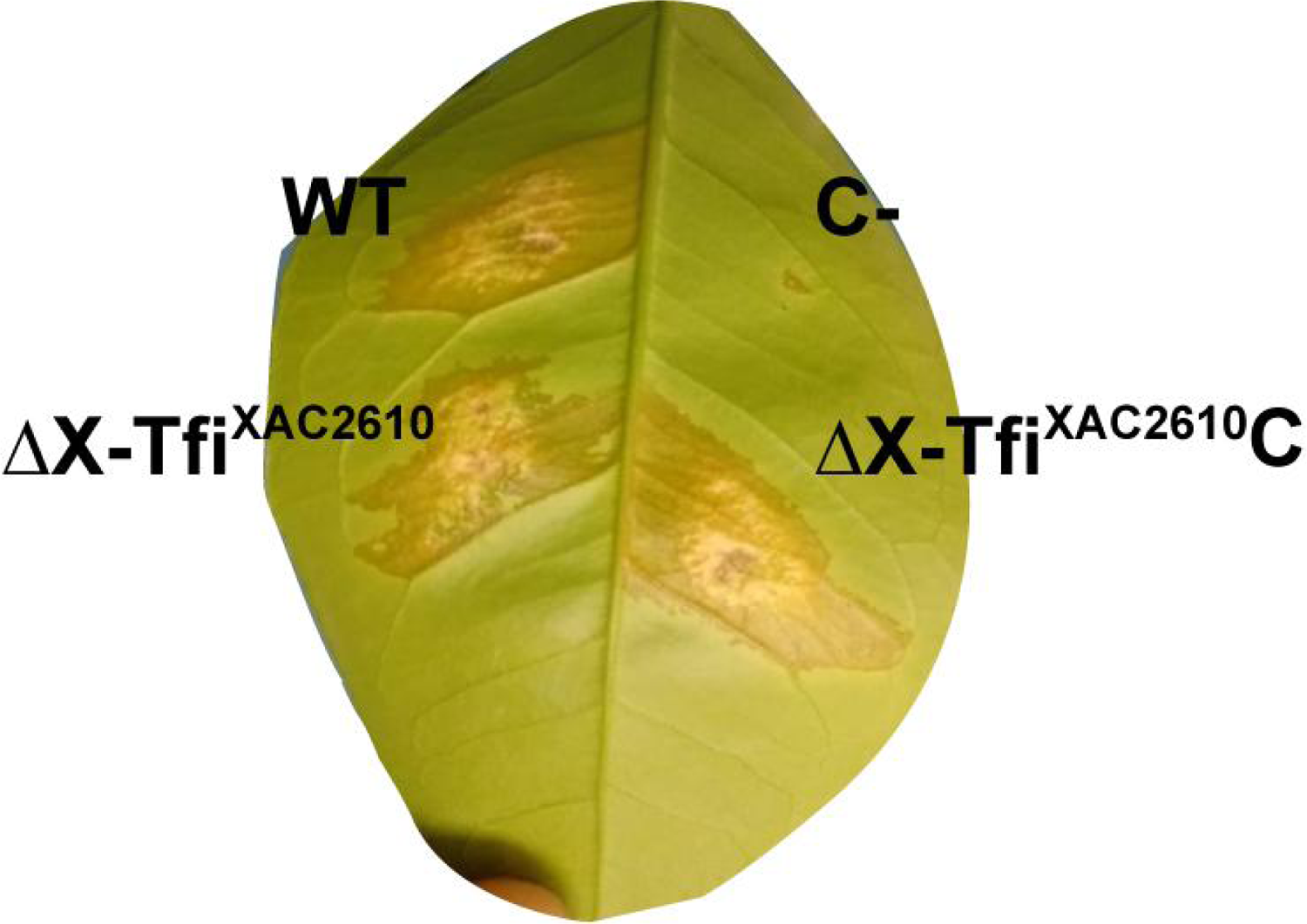
Citrus canker development assay on sweet orange leaves after 14 days of infection. *X. citri strains used: wild-type (WT),* ΔX-Tfi^XAC2610^, ΔX-Tfi^XAC2610^ complemented with pBRAXAC2610 vector (ΔX-Tfi^XAC2610^C). C^-^: mock infection with distilled water.

**Figure S7.**
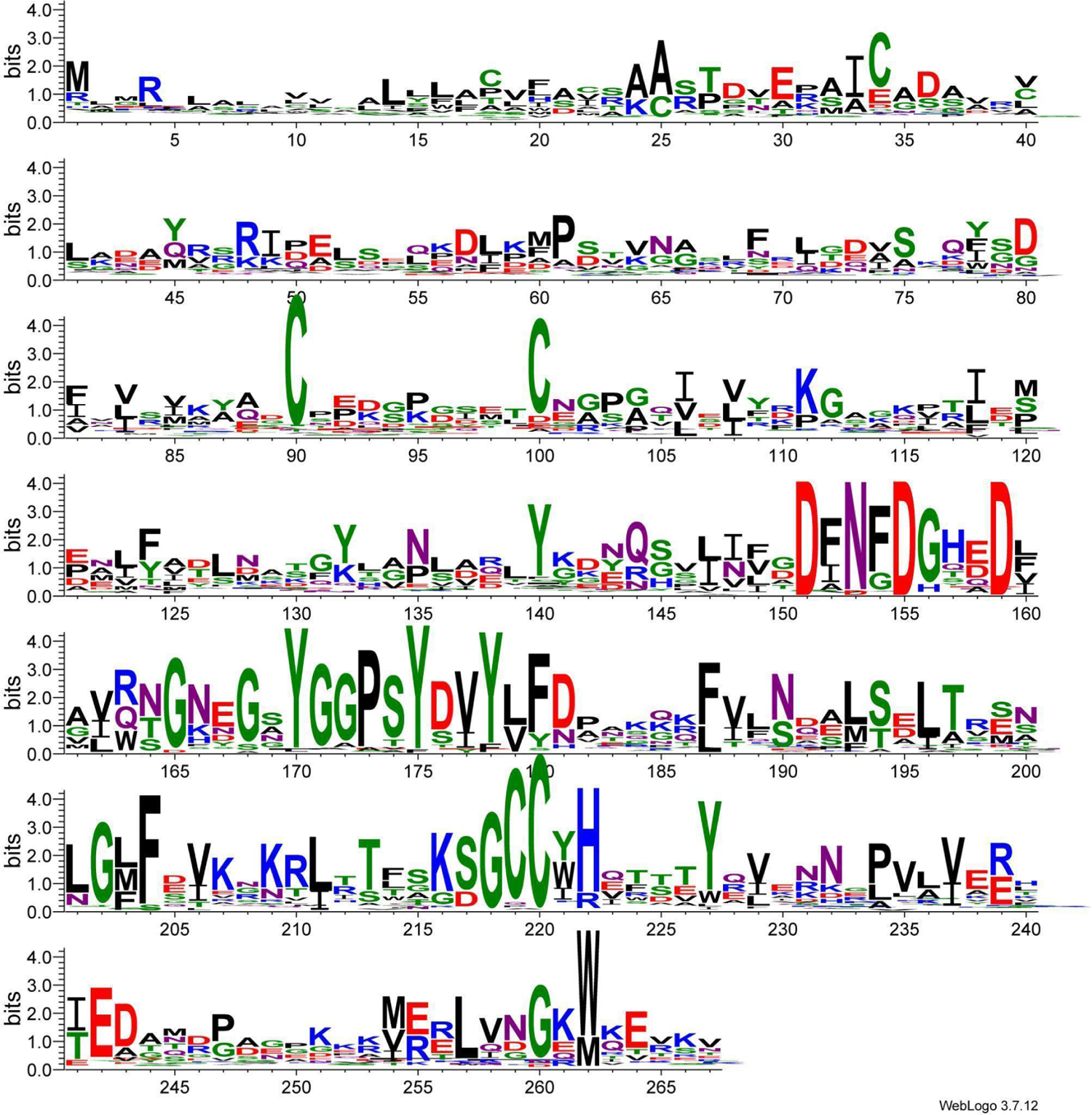
Sequence conservation profile of the multiple alignment of X-Tfi^XAC2610^ homologs. The sequences of 429 X-Tfi^XAC2610^ homologs (GenBank code AAM37459/ ref seq WP_011051709.1) were obtained as described in Material and Methods and are listed in **Supplementary File S1**. The sequence conservation profile was generated using the Weblogo3 server (Crooks et al. 2004). Stacking height indicates sequence conservation at each position while the symbol within the stack indicates the relative frequency of each amino acid in that position. The numbering below the profile corresponds to the amino acid sequence of X-Tfi^XAC2610^. The conserved motif from residues 151-159 corresponds to the Ca^2+^-binding loop observed in the X-Tfi^XAC2610^ crystal structure (Souza et al. 2015). The conserved tyrosine at position 170 is found within a loop predicted to insert into the active site of the cognate effector as described in the main text.

**Figure S8.**
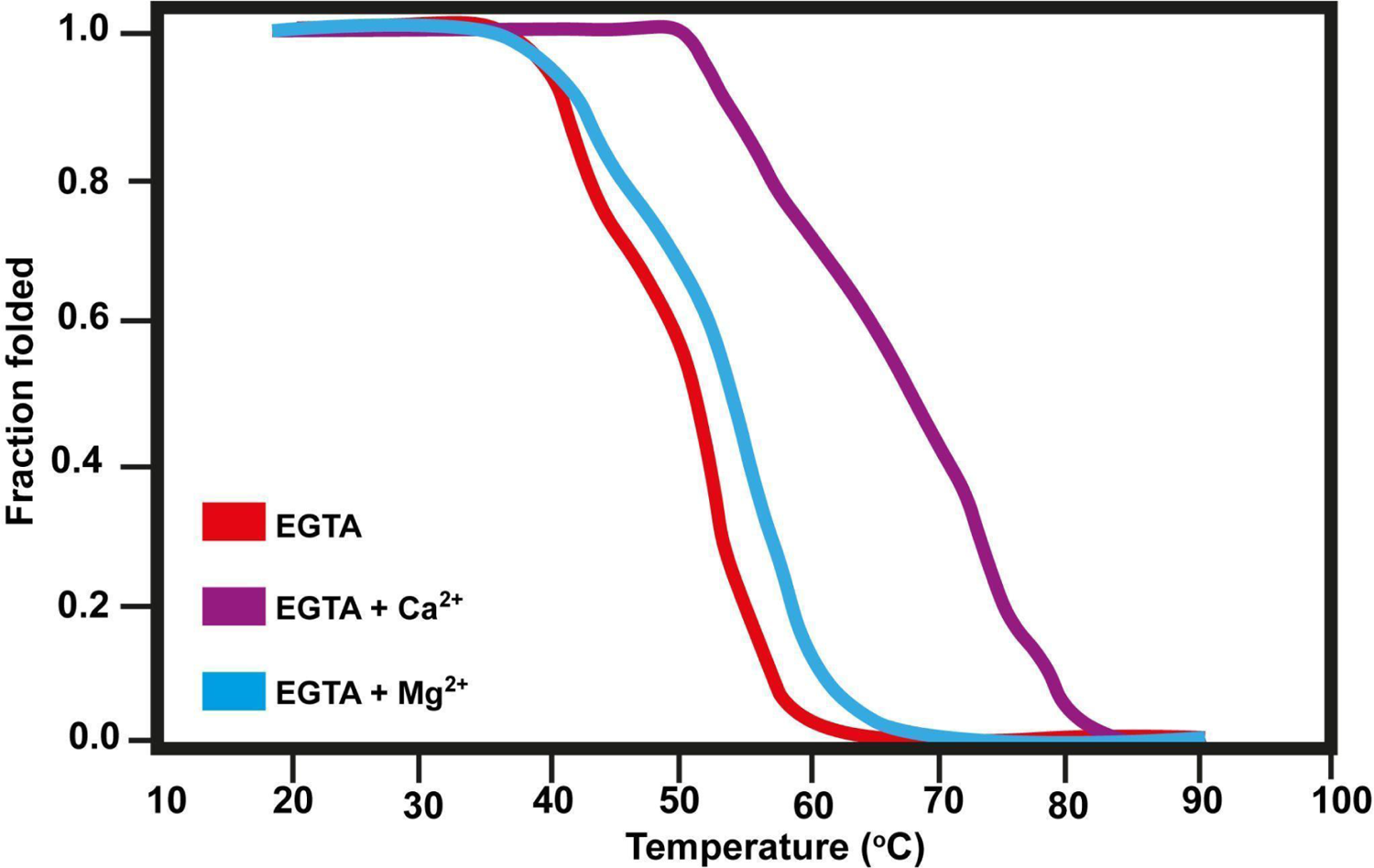
Ca^2+^ binding stabilizes X-Tfi^XAC2610^. Thermal denaturation experiments were conducted with X-Tfi^XAC2610^(55-267) (0.5 mM protein in 20 mM Tris-HCl (pH 7.5) and 50 mM NaCl) in the presence of EGTA (red), EGTA and Ca^2+^ (purple), EGTA and Mg^2+^ (cyan). EGTA and divalent metal salts (MgCl_2_ and CaCl_2_) were added with final concentrations of 0.5 mM and 0.75 mM, respectively. The fraction of folded protein was calculated by monitoring the intrinsic tryptophan fluorescence as described in the Materials and Methods (Pace and Scholtz 1997).

**Figure S9.**
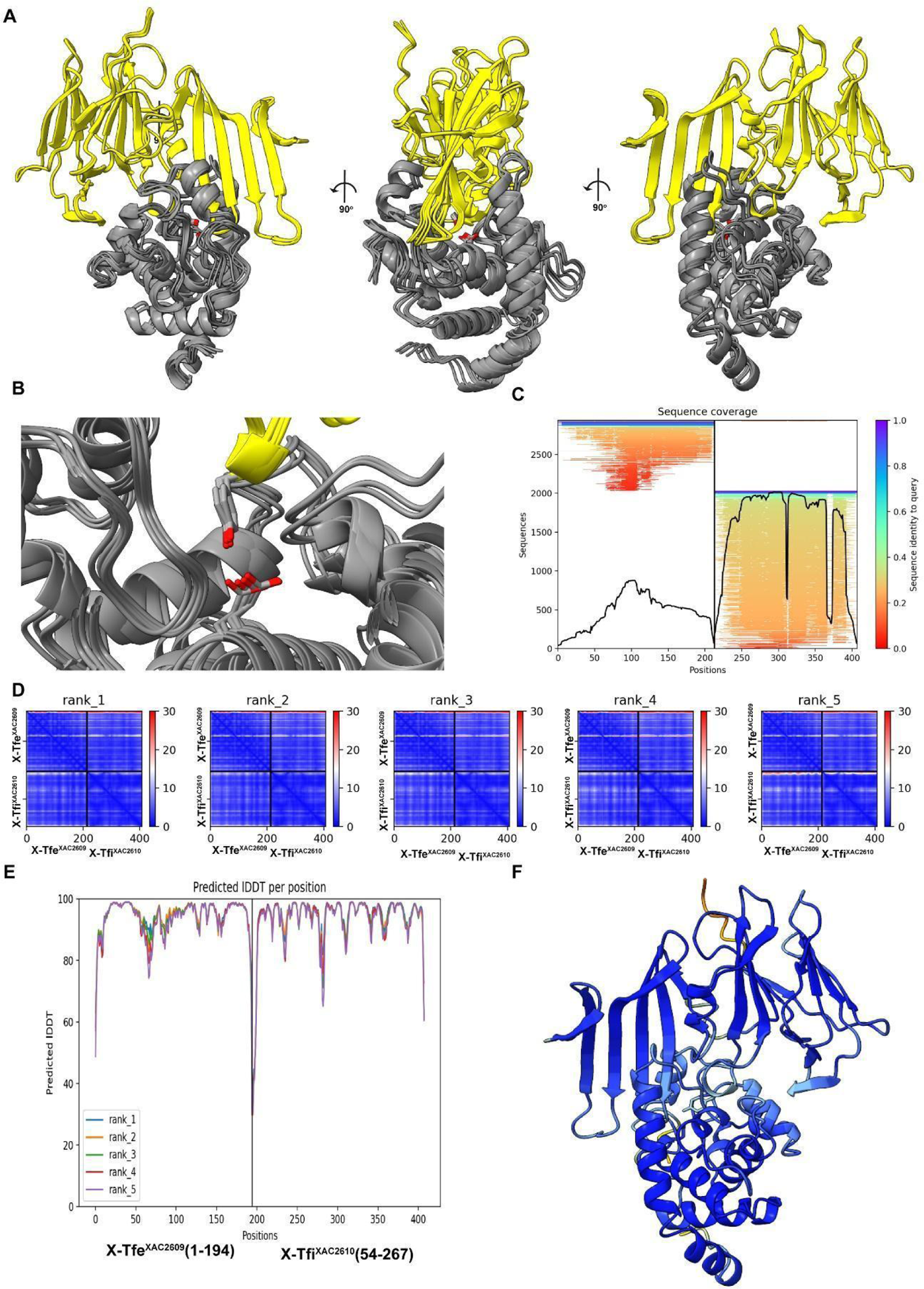
AlphaFold 2 models of the X-Tfe^XAC2609^-X-Tfi^XAC2610^ complex. **(A)** Superposition of 5 models of the X-Tfe^XAC2609^(1-194)-X-Tfi^XAC2610^(54-267) complex predicted by ColabFold-AlphaFold 2 (Mirdita et al. 2022; Varadi et al. 2022; Jumper et al. 2021). X-Tfi^XAC2610^(54-267) (yellow); X-Tfe^XAC2609^(1-194) (gray). **(B)** Interaction between the conserved Y170 of X-Tfi^XAC2610^ and E48 of X-Tfe^XAC2609^ from the ensemble shown in (**A**). Y170 and E48 are shown as stick models. **(C)** Multiple Sequence coverage of the X-Tfe^XAC2609^(1-194)-X-Tfi^XAC2610^(54-267) complex. **(D)** Predicted Aligned Error file (PAE) of 5 models of the X-Tfe^XAC2609^(1-194)-X-Tfi^XAC2610^(54-267) complex. **(E)** Predicted local Distance Difference Test (lDDT) score. **(F)** Final model of the X-Tfe^XAC2609^(1-194)-X-Tfi^XAC2610^(54-267) complex colored according pLDDT values.

**Figure S10.**
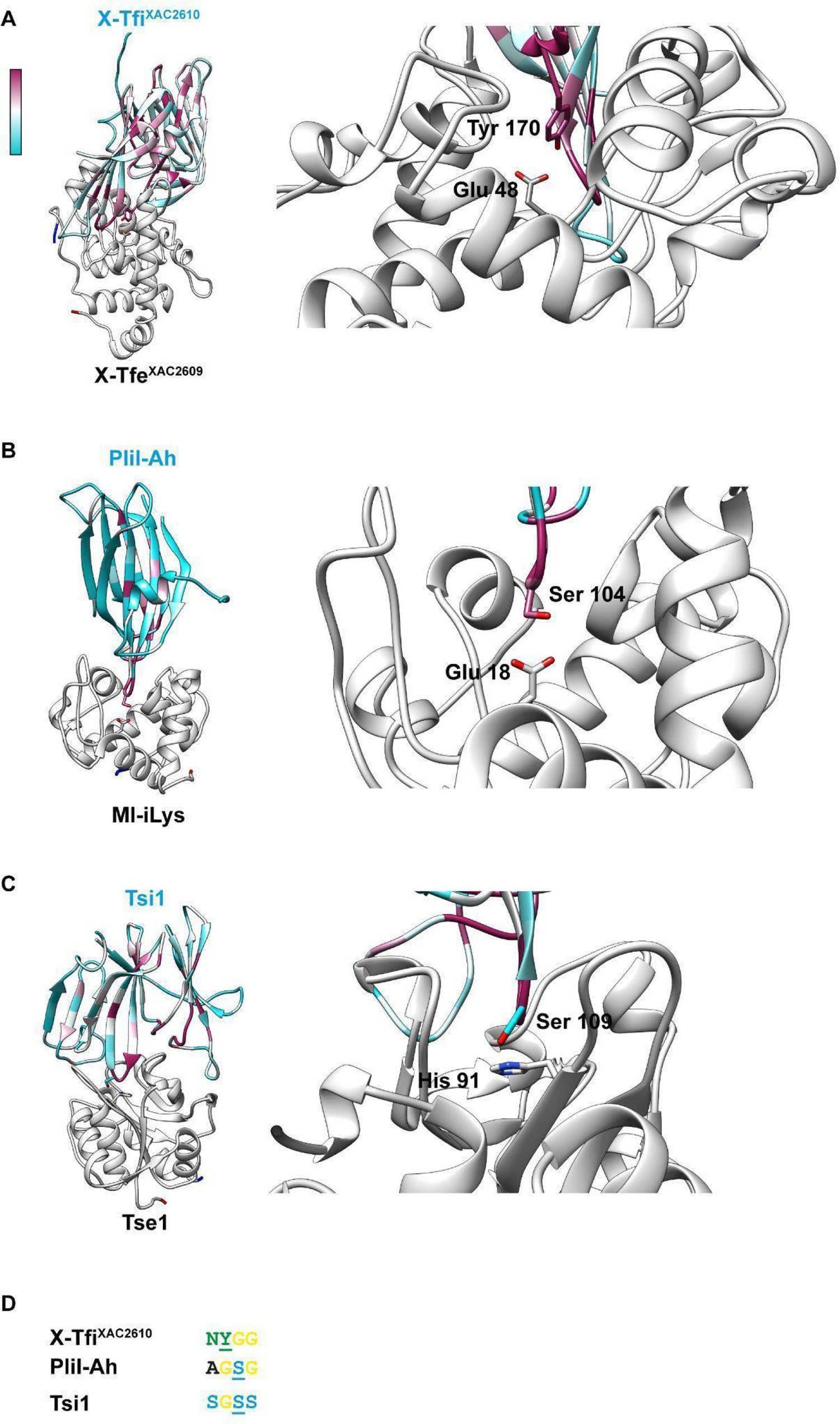
Comparison of the model of the X-Tfe^XAC2609^(1-194)-X-Tfi^XAC2610^(54-267) complex with the crystal structures of the PliI-Ah - i-type lysozyme and Tsi1-Tse1 complexes. A-C, *left panels*: Ribbon representation of the **(A)** X-Tfe^XAC2609^-X-Tfi^XAC2610^ model generated by Alpha-fold2. **(B)** the crystal structure of the periplasmic i-type lysozyme inhibitor from *Aeromonas hydrophila* (PliI-Ah) in complex with the i-type lysozyme from *Meretrix lusoria* (MI-iLys) (PDB 4PJ2; (Herreweghe et al. 2015) and **(C)** the crystal structure of Tsi1-Tse1 complex from (PDB 3VPJ) (Ding et al. 2012). In **A, B** and **C**, the inhibitors are colored according to the degree of conservation at each residue position in the corresponding family of homologs (lowest, cyan; highest, purple) and the cognate enzymes are colored in gray. *Right panels*: Zoom of the interaction interfaces showing the insertion of the inhibitory loops into the active sites of the enzymes. Stick models highlight the interactions between residues in the inhibitory loops (X-Tfi^XAC2610^ Y170, PliI-Ah S104 and Tsi1 S109) and the enzyme active sites (X-Tfe^XAC2609^ E48, MI-iLys E18, Tse1 H91). **D:** Sequences in the inhibitory loops of X-Tfi^XAC2610^, Plil-Ah and Tsi1 that interact directly with the catalytic site of the corresponding toxins. Underlined are X-Tfi^XAC2610^ Y170, PliI-Ah S104 and Tsi1 S109.

**Figure S11.**
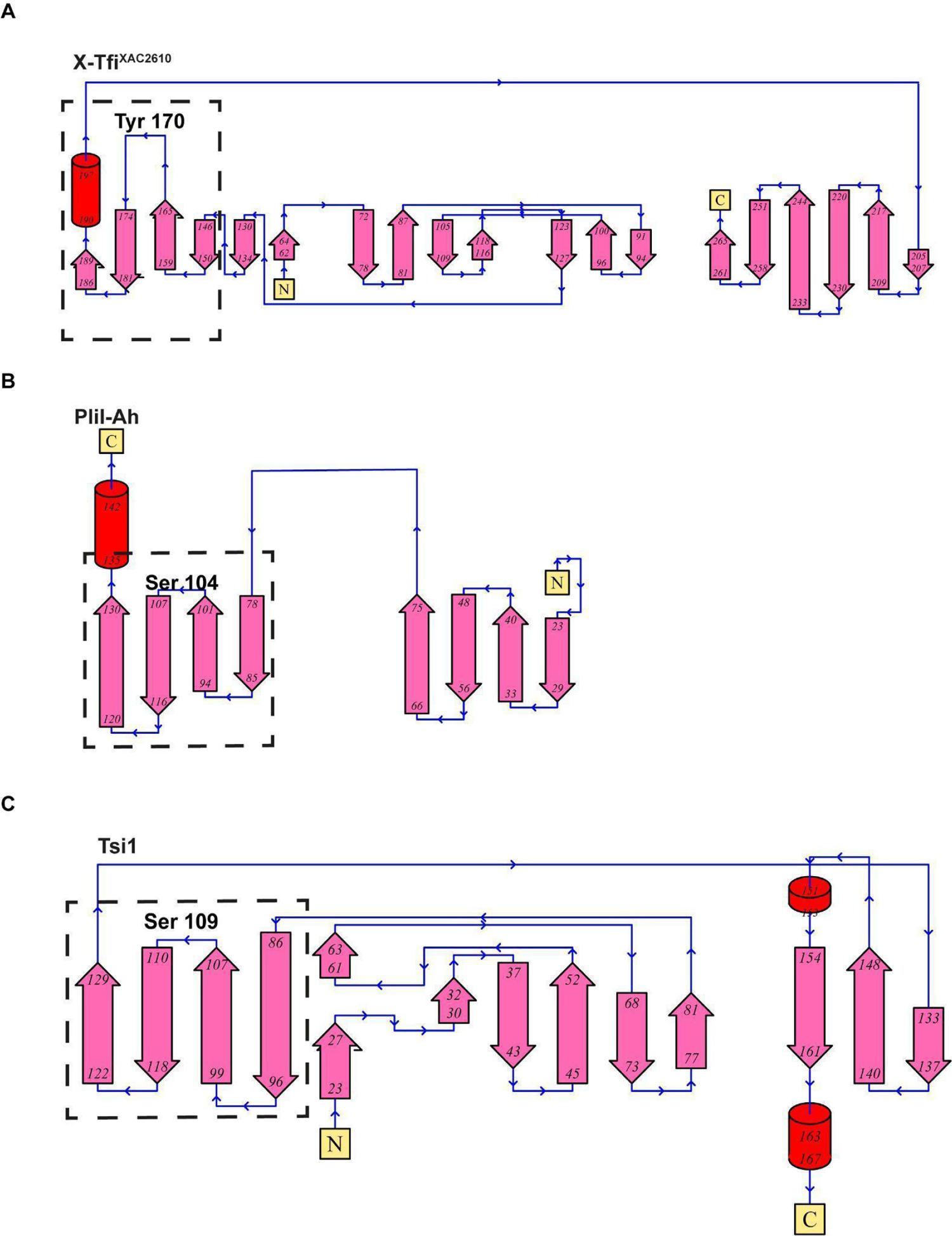
Protein topology diagrams of **(A)** X-Tfi^XAC2610^ (PDB 4QTQ), **(B)** Pli-Ah (PDB 4PJ2) and **(C)** Tsi1 (PDB 3VPJ). Diagrams were generated using the PDBsum server (http://www.ebi.ac.uk/thornton-srv/databases/cgi-bin/pdbsum/GetPage.pl?pdbcode=ind ex.html). Secondary-structure elements are indicated as red cylinders (α-helices) and pink arrows (β-strands). The dotted-squares highlight the common β-sheet found in the three immunity proteins containing a loop between the second and third β-strands that inserts into the active site of the cognate enzyme.

## REFERENCES

Alegria, Marcos C., Diorge P. Souza, Maxuel O. Andrade, Cassia Docena, Leticia Khater, Carlos H. I. Ramos, Ana C. R. da Silva, and Chuck S. Farah. 2005. “Identification of New Protein-Protein Interactions Involving the Products of the Chromosome- and Plasmid-Encoded Type IV Secretion Loci of the Phytopathogen Xanthomonas Axonopodis Pv. Citri.” Journal of Bacteriology 187 (7): 2315–25.

Alvarez-Martinez, Cristina E., and Peter J. Christie. 2009. “Biological Diversity of Prokaryotic Type IV Secretion Systems.” Microbiology and Molecular Biology Reviews: MMBR 73 (4): 775–808.

Aoki, Stephanie K., Rupinderjit Pamma, Aaron D. Hernday, Jessica E. Bickham, Bruce A. Braaten, and David A. Low. 2005. “Contact-Dependent Inhibition of Growth in Escherichia Coli.” Science 309 (5738): 1245–48.

Atmakuri, Krishnamohan, Eric Cascales, and Peter J. Christie. 2004. “Energetic Components VirD4, VirB11 and VirB4 Mediate Early DNA Transfer Reactions Required for Bacterial Type IV Secretion.” Molecular Microbiology 54 (5): 1199–1211.

Barnes, Christopher O., and Gary J. Pielak. 2011. “In-Cell Protein NMR and Protein Leakage.” Proteins 79 (2): 347–51.

Basler, M., and J. J. Mekalanos. 2012. “Type 6 Secretion Dynamics Within and Between Bacterial Cells.” Science. 10.1126/science.1222901.

Bayer-Santos, Ethel, William Cenens, Bruno Yasui Matsuyama, Gabriel Umaji Oka, Giancarlo Di Sessa, Izabel Del Valle Mininel, Tiago Lubiana Alves, and Chuck Shaker Farah. 2019. “The Opportunistic Pathogen Stenotrophomonas Maltophilia Utilizes a Type IV Secretion System for Interbacterial Killing.” PLoS Pathogens 15 (9): e1007651.

Benz, Juliane, and Anton Meinhart. 2014. “Antibacterial Effector/immunity Systems: It’s Just the Tip of the Iceberg.” Current Opinion in Microbiology 17 (February): 1–10.

Benz, Juliane, Christina Sendlmeier, Thomas R. M. Barends, and Anton Meinhart. 2012. “Structural Insights into the Effector – Immunity System Tse1 / Tsi1 from Pseudomonas Aeruginosa” 7 (7). 10.1371/journal.pone.0040453.

Bucher, Tabitha, Yaara Oppenheimer-Shaanan, Alon Savidor, Zohar Bloom-Ackermann, and Ilana Kolodkin-Gal. 2015. “Disturbance of the Bacterial Cell Wall Specifically Interferes with Biofilm Formation.” Environmental Microbiology Reports 7 (6): 990–1004.

Callewaert, Lien, and Chris W. Michiels. 2010. “Lysozymes in the Animal Kingdom.” Journal of Biosciences 35 (1): 127–60.

Callewaert, Lien, Joris M. Van Herreweghe, Lise Vanderkelen, Seppe Leysen, Arnout Voet, and Chris W. Michiels. 2012. “Guards of the Great Wall: Bacterial Lysozyme Inhibitors.” Trends in Microbiology 20 (10): 501–10.

Cao, Zhenping, M. Guillermina Casabona, Holger Kneuper, James D. Chalmers, and Tracy Palmer. 2016. “The Type VII Secretion System of Staphylococcus Aureus Secretes a Nuclease Toxin That Targets Competitor Bacteria.” Nature Microbiology 2 (October): 16183.

Cappelletti, Paola A., Rafael Freitas dos Santos, Alexandre M. do Amaral, Rafael Augusto Homem, Thaís dos Santos Souza, Marcos A. Machado, and Chuck S. Farah. 2011. “Structure-Function Analysis of the HrpB2-HrcU Interaction in the Xanthomonas Citri Type III Secretion System.” PloS One 6 (3): e17614.

Cascales, Eric, and Peter J. Christie. 2004a. “Definition of a Bacterial Type IV Secretion Pathway for a DNA Substrate.” Science 304 (5674): 1170–73.

Cascales, Eric, and Peter J. Christie. 2004b. “Agrobacterium VirB10, an ATP Energy Sensor Required for Type IV Secretion.” Proceedings of the National Academy of Sciences of the United States of America 101 (49): 17228–33.

Cernooka, Elina, Janis Rumnieks, Nikita Zrelovs, Kaspars Tars, and Andris Kazaks. 2022. “Diversity of the Lysozyme Fold: Structure of the Catalytic Domain from an Unusual Endolysin Encoded by Phage Enc34.” Scientific Reports 12 (1): 5005.

Chandran, Vidya, Rémi Fronzes, Stéphane Duquerroy, Nora Cronin, Gabriel Waksman, Jorge Navaza, Gabriel Waksman, and Jorge Navaza. 2009. “Structure of the Outer Membrane Complex of a Type IV Secretion System.” Nature 462 (7276): 1011–15.

Christie, Peter J., Neal Whitaker, and Christian González-Rivera. 2014. “Mechanism and Structure of the Bacterial Type IV Secretion Systems.” Biochimica et Biophysica Acta 1843 (8): 1578–91.

Cianfanelli, Francesca R., Laura Monlezun, and Sarah J. Coulthurst. 2016. “Aim, Load, Fire: The Type VI Secretion System, a Bacterial Nanoweapon.” Trends in Microbiology 24 (1): 51–62.

Costa, Tiago R. D., Catarina Felisberto-Rodrigues, Amit Meir, Marie S. Prevost, Adam Redzej, Martina Trokter, and Gabriel Waksman. 2015. “Secretion Systems in Gram-Negative Bacteria: Structural and Mechanistic Insights.” Nature Reviews. Microbiology 13 (6): 343–59.

Crooks, Gavin E., Gary Hon, John-Marc Chandonia, and Steven E. Brenner. 2004. “WebLogo: A Sequence Logo Generator.” Genome Research 14 (6): 1188–90.

Ding, Jingjin, Wei Wang, Han Feng, Ying Zhang, and Da-Cheng Wang. 2012. “Structural Insights into the Pseudomonas Aeruginosa Type VI Virulence Effector Tse1 Bacteriolysis and Self-Protection Mechanisms.” The Journal of Biological Chemistry 287 (32): 26911–20.

Dong, Tao G., Brian T. Ho, Deborah R. Yoder-Himes, and John J. Mekalanos. 2013. “Identification of T6SS-Dependent Effector and Immunity Proteins by Tn-Seq in Vibrio Cholerae.” Proceedings of the National Academy of Sciences of the United States of America 110 (7): 2623–28.

Dooley, Erin E. 2004. “National Center for Biotechnology Information.” Environmental Health Perspectives 112 (12): A674.

Dunger, German, Cristiane R. Guzzo, Maxuel O. Andrade, Jeffrey B. Jones, and Chuck S. Farah. 2014. “Xanthomonas Citri Subsp. Citri Type IV Pilus Is Required for Twitching Motility, Biofilm Development, and Adherence.” Molecular Plant-Microbe Interactions: MPMI 27 (10): 1132–47.

Dunger, German, Edgar Llontop, Cristiane R. Guzzo, and Chuck S. Farah. 2016. “ScienceDirect The Xanthomonas Type IV Pilus.” Current Opinion in Microbiology 30 (April): 88–97.

Edgar, Robert C. 2004. “MUSCLE: Multiple Sequence Alignment with High Accuracy and High Throughput.” Nucleic Acids Research 32 (5): 1792–97.

Fleming, Alexander, and Almroth Edward Wright. 1922. “On a Remarkable Bacteriolytic Element Found in Tissues and Secretions.” *Proceedings of the Royal Society of London. Series B*, Containing Papers of a Biological Character. Royal Society 93 (653): 306–17.

Fronzes, Rémi, Peter J. Christie, and Gabriel Waksman. 2009. “The Structural Biology of Type IV Secretion Systems.” Nature Reviews. Microbiology 7 (10): 703–14.

García-Bayona, Leonor, Monica S. Guo, and Michael T. Laub. 2017. “Contact-Dependent Killing by Caulobacter Crescentus via Cell Surface-Associated, Glycine Zipper Proteins.” eLife 6 (March). 10.7554/eLife.24869.

Herreweghe, Van, Seppe Leysen, Joris M. Van Herreweghe, Kazunari Yoneda, Makoto Ogata, Taichi Usui, Tomohiro Araki, Christiaan W. Michiels, and Sergei V. Strelkov. 2015. “The Structure of the Proteinaceous Inhibitor PliI from Aeromonas Hydrophila in Complex with Its Target Lysozyme.” *Acta Crystallographica. Section D*, Biological Crystallography 71 (Pt 2): 344–51.

Ho, Brian T., Yang Fu, Tao G. Dong, and John J. Mekalanos. 2017. “*Vibrio Cholerae* Type 6 Secretion System Effector Trafficking in Target Bacterial Cells.” Proceedings of the National Academy of Sciences of the United States of America 114 (35): 9427–32.

Hood, Rachel D., Pragya Singh, Fosheng Hsu, Tüzün Güvener, Mike A. Carl, Rex R. S. Trinidad, Julie M. Silverman, et al. 2010. “A Type VI Secretion System of Pseudomonas Aeruginosa Targets a Toxin to Bacteria.” Cell Host & Microbe 7 (1): 25–37.

Ilangovan, Aravindan, Sarah Connery, and Gabriel Waksman. 2015. “Structural Biology of the Gram-Negative Bacterial Conjugation Systems.” Trends in Microbiology 23 (5): 301–10.

Iwata, Taketoshi, Ayako Watanabe, Masahiro Kusumoto, and Masato Akiba. 2016. “Peptidoglycan Acetylation of Campylobacter Jejuni Is Essential for Maintaining Cell Wall Integrity and Colonization in Chicken Intestine.” Applied and Environmental Microbiology 82 (20): 6284–90.

Jumper, John, Richard Evans, Alexander Pritzel, Tim Green, Michael Figurnov, Olaf Ronneberger, Kathryn Tunyasuvunakool, et al. 2021. “Highly Accurate Protein Structure Prediction with AlphaFold.” Nature 596 (7873): 583–89.

Jurėnas, Dukas, and Laure Journet. 2021. “Activity, Delivery, and Diversity of Type VI Secretion Effectors.” Molecular Microbiology 115 (3): 383–94.

Kobayashi, Kazuo. 2021. “Diverse LXG Toxin and Antitoxin Systems Specifically Mediate Intraspecies Competition in Bacillus Subtilis Biofilms.” PLoS Genetics 17 (7): e1009682.

Li, Mo, Isolde Le Trong, Mike A. Carl, Eric T. Larson, Seemay Chou, Justin A. De Leon, Simon L. Dove, Ronald E. Stenkamp, and Joseph D. Mougous. 2012. “Structural Basis for Type VI Secretion Effector Recognition by a Cognate Immunity Protein.” PLoS Pathogens 8 (4): e1002613.

Li, Yang Grace, and Peter J. Christie. 2018. “The Agrobacterium VirB/VirD4 T4SS: Mechanism and Architecture Defined Through In Vivo Mutagenesis and Chimeric Systems.” Current Topics in Microbiology and Immunology 418: 233–60.

Lowe, G., G. Sheppard, M. L. Sinnott, and A. Williams. 1967. “Lysozyme-Catalysed Hydrolysis of Some Beta-Aryl Di-N-Acetylchitobiosides.” Biochemical Journal 104 (3): 893–99.

Macé, Kévin, Abhinav K. Vadakkepat, Adam Redzej, Natalya Lukoyanova, Clasien Oomen, Nathalie Braun, Marta Ukleja, et al. 2022. “Cryo-EM Structure of a Type IV Secretion System.” Nature 607 (7917): 191–96.

MacIntyre, Dana L., Sarah T. Miyata, Maya Kitaoka, and Stefan Pukatzki. 2010. “The Vibrio Cholerae Type VI Secretion System Displays Antimicrobial Properties.” Proceedings of the National Academy of Sciences of the United States of America 107 (45): 19520–24.

Metcalf, Jason A., Lisa J. Funkhouser-Jones, Kristen Brileya, Anna-Louise Reysenbach, and Seth R. Bordenstein. 2014. “Antibacterial Gene Transfer across the Tree of Life.” eLife 3 (November). 10.7554/eLife.04266.

Mirdita, Milot, Konstantin Schütze, Yoshitaka Moriwaki, Lim Heo, Sergey Ovchinnikov, and Martin Steinegger. 2022. “ColabFold: Making Protein Folding Accessible to All.” Nature Methods 19 (6): 679–82.

Murdoch, Sarah L., Katharina Trunk, Grant English, Maximilian J. Fritsch, Ehsan Pourkarimi, and Sarah J. Coulthurst. 2011. “The Opportunistic Pathogen Serratia Marcescens Utilizes Type VI Secretion to Target Bacterial Competitors.” Journal of Bacteriology 193 (21): 6057–69.

Navarro, Paula P., Andrea Vettiger, Virly Y. Ananda, Paula Montero Llopis, Christoph Allolio, Thomas G. Bernhardt, and Luke H. Chao. 2022. “Cell Wall Synthesis and Remodelling Dynamics Determine Division Site Architecture and Cell Shape in Escherichia Coli.” Nature Microbiology. 10.1038/s41564-022-01210-z.

Oka, Gabriel U., Diorge P. Souza, William Cenens, Bruno Y. Matsuyama, Marcus V. C. Cardoso, Luciana C. Oliveira, Filipe da Silva Lima, et al. 2022. “Structural Basis for Effector Recognition by an Antibacterial Type IV Secretion System.” Proceedings of the National Academy of Sciences of the United States of America 119 (1). 10.1073/pnas.2112529119.

Oliveira, Luciana Coutinho, Diorge Paulo Souza, Gabriel Umaji Oka, Filipe da Silva Lima, Ronaldo Junio Oliveira, Denize Cristina Favaro, Hans Wienk, Rolf Boelens, Chuck Shaker Farah, and Roberto Kopke Salinas. 2016. “VirB7 and VirB9 Interactions Are Required for the Assembly and Antibacterial Activity of a Type IV Secretion System.” Structure 24 (10): 1707–18.

Pace, C. Nick, and J. Martin Scholtz. 1997. “Measuring the Conformational Stability of a Protein.” Protein Structure: A Practical Approach 2: 299–321.

Pettersen, Eric F., Thomas D. Goddard, Conrad C. Huang, Elaine C. Meng, Gregory S. Couch, Tristan I. Croll, John H. Morris, and Thomas E. Ferrin. 2021. “UCSF ChimeraX: Structure Visualization for Researchers, Educators, and Developers.” Protein Science: A Publication of the Protein Society 30 (1): 70–82.

Russell, Alistair B., Rachel D. Hood, Nhat Khai Bui, Michele LeRoux, Waldemar Vollmer, and Joseph D. Mougous. 2011. “Type VI Secretion Delivers Bacteriolytic Effectors to Target Cells.” Nature 475 (7356): 343–47.

Russell, Alistair B., S. Brook Peterson, and Joseph D. Mougous. 2014. “Type VI Secretion System Effectors: Poisons with a Purpose.” Nature Reviews. Microbiology 12 (2): 137–48.

Russell, Alistair B., Pragya Singh, Mitchell Brittnacher, Nhat Khai Bui, Rachel D. Hood, Mike A. Carl, Danielle M. Agnello, et al. 2012. “A Widespread Bacterial Type VI Secretion Effector Superfamily Identified Using a Heuristic Approach.” Cell Host & Microbe 11 (5): 538–49.

Sana, Thibault G., Nicolas Flaugnatti, Kyler A. Lugo, Lilian H. Lam, Amanda Jacobson, Virginie Baylot, Eric Durand, Laure Journet, Eric Cascales, and Denise M. Monack. 2016. “Salmonella Typhimurium Utilizes a T6SS-Mediated Antibacterial Weapon to Establish in the Host Gut.” Proceedings of the National Academy of Sciences of the United States of America 113 (34): E5044–51.

Schindelin, Johannes, Ignacio Arganda-Carreras, Erwin Frise, Verena Kaynig, Mark Longair, Tobias Pietzsch, Stephan Preibisch, et al. 2012. “Fiji: An Open-Source Platform for Biological-Image Analysis.” Nature Methods 9 (7): 676–82.

Schwarz, Sandra, T. Eoin West, Frédéric Boyer, Wen-Chi Chiang, Mike A. Carl, Rachel D. Hood, Laurence Rohmer, Tim Tolker-Nielsen, Shawn J. Skerrett, and Joseph D. Mougous. 2010. “Burkholderia Type VI Secretion Systems Have Distinct Roles in Eukaryotic and Bacterial Cell Interactions.” PLoS Pathogens 6 (8): e1001068.

Sena-Vélez, Marta, Cristina Redondo, James H. Graham, and Jaime Cubero. 2016. “Presence of Extracellular DNA during Biofilm Formation by Xanthomonas Citri Subsp. Citri Strains with Different Host Range.” PloS One 11 (6): 1–17.

Sgro, Germán G., Tiago R. D. Costa, William Cenens, Diorge P. Souza, Alexandre Cassago, Luciana Coutinho de Oliveira, Roberto K. Salinas, Rodrigo V. Portugal, Chuck S. Farah, and Gabriel Waksman. 2018. “Cryo-EM Structure of the Bacteria-Killing Type IV Secretion System Core Complex from Xanthomonas Citri.” Nature Microbiology 3 (12): 1429–40.

Sgro, Germán G., Gabriel U. Oka, Diorge P. Souza, William Cenens, Ethel Bayer-Santos, Bruno Y. Matsuyama, Natalia F. Bueno, et al. 2019. “Bacteria-Killing Type IV Secretion Systems.” Frontiers in Microbiology 10 (May): 1078.

Sheedlo, Michael J., Melanie D. Ohi, D. Borden Lacy, and Timothy L. Cover. 2022. “Molecular Architecture of Bacterial Type IV Secretion Systems.” PLoS Pathogens 18 (8): e1010720.

Silhavy, Thomas J., Daniel Kahne, and Suzanne Walker. 2010. “The Bacterial Cell Envelope.” Cold Spring Harbor Perspectives in Biology 2 (5): a000414.

Silva, A. C. R. da, J. A. Ferro, F. C. Reinach, C. S. Farah, L. R. Furlan, R. B. Quaggio, C. B. Monteiro-Vitorello, et al. 2002. “Comparison of the Genomes of Two Xanthomonas Pathogens with Differing Host Specificities.” Nature 417 (6887): 459–63.

Souza, Diorge P., Maxuel O. Andrade, Cristina E. Alvarez-Martinez, Guilherme M. Arantes, Chuck S. Farah, and Roberto K. Salinas. 2011. “A Component of the Xanthomonadaceae Type IV Secretion System Combines a VirB7 Motif with a N0 Domain Found in Outer Membrane Transport Proteins.” PLoS Pathogens 7 (5): e1002031.

Souza, Diorge P., Gabriel U. Oka, Cristina E. Alvarez-Martinez, Alexandre W. Bisson-Filho, German Dunger, Lise Hobeika, Nayara S. Cavalcante, et al. 2015. “Bacterial Killing via a Type IV Secretion System.” Nature Communications 6 (March): 6453.

Tassinari, Matteo, Thierry Doan, Marco Bellinzoni, Maïalene Chabalier, Mathilde Ben-Assaya, Mariano Martinez, Quentin Gaday, et al. 2022. “The Antibacterial Type VII Secretion System of Bacillus Subtilis: Structure and Interactions of the Pseudokinase YukC/EssB.” *mBio*, September, e0013422.

Teufel, Felix, José Juan Almagro Armenteros, Alexander Rosenberg Johansen, Magnús Halldór Gíslason, Silas Irby Pihl, Konstantinos D. Tsirigos, Ole Winther, Søren Brunak, Gunnar von Heijne, and Henrik Nielsen. 2022. “SignalP 6.0 Predicts All Five Types of Signal Peptides Using Protein Language Models.” Nature Biotechnology. 10.1038/s41587-021-01156-3.

Varadi, Mihaly, Stephen Anyango, Mandar Deshpande, Sreenath Nair, Cindy Natassia, Galabina Yordanova, David Yuan, et al. 2022. “AlphaFold Protein Structure Database: Massively Expanding the Structural Coverage of Protein-Sequence Space with High-Accuracy Models.” Nucleic Acids Research 50 (D1): D439–44.

Vocadlo, D. J., G. J. Davies, R. Laine, and S. G. Withers. 2001. “Catalysis by Hen Egg-White Lysozyme Proceeds via a Covalent Intermediate.” Nature 412 (6849): 835–38.

Watanabe, I., and E. Yamada. 1983. “The Fine Structure of Lamellated Nerve Endings Found in the Rat Gingiva.” Archivum Histologicum Japonicum = Nihon Soshikigaku Kiroku 46 (2): 173–82.

Whitney, John C., Seemay Chou, Alistair B. Russell, Jacob Biboy, Taylor E. Gardiner, Michael A. Ferrin, Mitchell Brittnacher, Waldemar Vollmer, and Joseph D. Mougous. 2013. “Identification, Structure, and Function of a Novel Type VI Secretion Peptidoglycan Glycoside Hydrolase Effector-Immunity Pair.” The Journal of Biological Chemistry 288 (37): 26616–24.

Whitney, John C., Dennis Quentin, Shin Sawai, Michele LeRoux, Brittany N. Harding, Hannah E. Ledvina, Bao Q. Tran, et al. 2015. “An Interbacterial NAD(P)(+) Glycohydrolase Toxin Requires Elongation Factor Tu for Delivery to Target Cells.” Cell 163 (3): 607–19.

